# Adenosine deaminase co-immunization reverses age-associated immunosenescence by restoring germinal center T follicular helper cell function

**DOI:** 10.64898/2026.06.04.730208

**Authors:** Emily N. Bitsko, David R. Joyner, Jeffrey R. Maslanka, Kyra Woloszczuk, Arden O. Edgerton, Matthew R. Bell, Christian Sell, Mohamad-Gabriel Alameh, Joris Beld, Elias K. Haddad, Michael C. Abt, Michele A. Kutzler

## Abstract

Older adults experience disproportionate morbidity and mortality from *Clostridioides difficile* infection (CDI); however, existing toxoid- and receptor binding domain (RBD)-based vaccines elicit suboptimal protection in aged hosts due to the age-associated defects in CD4^+^ T cell function, T follicular helper (T_FH_) cell activation, and antibody quality. We evaluated whether adenosine deaminase (ADA), an enzymatic immune modulator that degrades immunosuppressive adenosine, and improves GC T_FH_ differentiation and survival, could reverse these age-related impairments when co-delivered with DNA vaccine plasmids targeting toxin A and B RBDs (pRBD). In aged mice, pRBD vaccination alone produced markedly reduced toxin-specific effector/memory CD4^+^ T cells, diminished T_FH_ activation, and poor toxin A neutralization compared to vaccinated young mice. Co-immunization with plasmid-encoded adenosine deaminase-1 (pADA) restored toxin-specific CD4^+^ T cell generation and cytokine production, activation-induced marker (AIM) T_FH_ responses, and antibody-mediate toxin neutralization to levels comparable to young adults. Mechanistically, pADA co-immunization was associated with the reduction of CXCR4 on germinal center (GC) T_FH_ cells—an age-related defect linked to impaired GC positioning and diminished B cell help—suggesting that ADA improves humoral quality by correcting GC T_FH_ mislocalization. These immune enhancements corresponded with improved clinical outcomes in morbidity, mortality, and weight-loss following *C. difficile* spore challenge of aged mice. Finally, pADA significantly reduced adenosine levels in aged lymph nodes, implicating a potential enzymatic-based regulation of GC immunosenescence. Together, these findings identify ADA as a metabolic adjuvant capable of reversing key features of vaccine immunosenescence and highlight adenosine dependent CXCR4 regulation as a tractable axis for improving vaccine efficacy in older populations.

## Introduction

The elderly population (>65 years) is a rapidly growing at-risk group accounting for 10% of the world’s population and is projected to double by 2050(1). Reduced immune function and increased susceptibility to infectious diseases are primary drivers of morbidity and mortality in the elderly(2, 3). Vaccines therefore represent a critical strategy for inducing robust antigen-specific immunity that enables protective immunological recall upon subsequent infection. Unfortunately, vaccine-induced responses are often reduced and ineffective in the elderly due to declining immune function(4, 5). Specifically, impaired germinal center (GC) formation(6–8), T follicular helper cell (T_FH_) development and activation(9, 10), and diminished antigen-specific antibody production(11–13) are key mediators of failed vaccine-induced protection in the elderly. Aside from increasing immunization dose, the inclusion of adjuvants is a widely used approach to enhance vaccination responses in aged individuals. Despite being a crucial component of vaccines for the elderly, pre-clinical investigations of adjuvants in aged animal models are limited.

Among the pathogens that pose significant risk to the elderly, *Clostridioides difficile* is of particular importance due to the absence of a clinically available vaccine(14–16). *C. difficile* is a gram positive, spore-forming anaerobe and opportunistic pathogen that accounts for >50% of all hospital-acquired enteric infections, with elderly individuals disproportionately affected by *C. difficile* infection (CDI) and recurrence(17). Toxins A and B act as the primary virulence factors mediating *C. difficile* associated disease (CDAD)(18), with the receptor binding domains (RBDs) critical for initiating attachment and internalization into host cells. The formation of neutralizing antibody responses against *C. difficile* toxins has been implicated as a significant correlate of protection(19–26), further reinforced by the observed efficacy of Bezlotoxumab, a monoclonal antibody targeting toxin B RBD(27–29). In fact, toxoid and RBD subunit vaccine candidates sponsored by Pfizer(30), Sanofi(31), Valneva(32), and GSK(33) have progressed to clinical trials, but failed to reach their primary efficacy endpoints in adults and the elderly, suggesting inclusion of additional antigens(34) or novel adjuvants(35) may be necessary in future approaches(36). Although Pfizer’s detoxified toxoid vaccine did not meet its primary phase 3 efficacy endpoint, it reduced CDI duration and medically attended CDI in secondary analyses, highlighting that response quality—not merely antibody magnitude—is critical in older adults(30). Likewise, recent investigations of novel mucosal and mRNA-LNP vaccines comprised of inactivated toxin and non-toxin antigens demonstrated the influence of administration route on desired vaccine efficacy(34, 37). Considering the reported aging-related deficits in toxin-specific antibody production during CDI(38), we postulated that use of *C. difficile* vaccination and infection models would provide significant insight into adjuvant-mediated bolstering of aged humoral responses.

We have previously reported on the adjuvant-like properties of adenosine deaminase (ADA) in human leukocytes and murine vaccination models(39–41). ADA is a key enzyme required for the irreversible deamination of adenosine, an immunosuppressive nucleoside, into inosine. Tardif et al. established the role of ADA in delineating functional T_FH_ programming by altering the cytokine milieu, CD26 expression, and adenosine receptor signaling, suggesting ADA could exert adjuvant-like properties(41). In prior work, we demonstrated the propensity of ADA to function as an immune-modulator when co-administered with vaccines in a dose-sparing manner. Specifically, ADA was shown to enhance cytokine production promoting T_FH_ differentiation and survival, antigen-specific CD4/CD8 T cell activation and cytokine production, antibody titers, and antibody neutralization at low antigen doses using young murine vaccination models(39–42). However, the potential for ADA to restore impaired vaccine responses in aging requires further investigation. Activated T cells from aged individuals have been reported to produce higher levels of immunosuppressive adenosine(43), potentially linked to the age-associated expansion of CD39+ CD4^+^ T cells(44). Coupled with recent findings of age-induced CXCR4 upregulation on GC-T_FH_ as a driver of cellular mislocalization, diminished B cell help, and poor GC formation following vaccination(10, 45), we hypothesized the pro-T_FH_ and enzymatic activity of ADA could mechanistically restore these defects in an aged murine model.

Recent work on ADA as a molecular immune modulator has been primarily limited to low-dose vaccination studies in young murine models and does not mechanistically address how ADA could resolve vaccine immunosenescence in aging. Our development of a plasmid-encoded toxin RBD DNA vaccine(46) enables identification of adjuvant-mediated enhancements in vaccine efficacy. Here, we used a prime-boost strategy in young and aged mice to evaluate toxin-specific responses among major contributors to vaccine-elicited protection including CD4^+^ T effector/memory (T_EM_) cells, GC-T_FH_, and humoral immunity. We also apply a *C. difficile* challenge model that recapitulates age-related susceptibility to CDI(47, 48) to evaluate protection measured by morbidity, weight-loss, and mortality. pADA co-immunization restored toxin-specific CD4^+^ T cell generation and cytokine production, GC-T_FH_ activation, and antibody-mediated toxin neutralization. These improvements were accompanied by reduced CXCR4 expression on aged GC T_FH_, which in turn was collectively associated with reduced morbidity and mortality following lethal CD196 spore challenge. This work establishes that aged mice exhibit traditional markers of immunosenescence with our DNA vaccine and supports the use of ADA as a novel vaccine adjuvant for the elderly.

## Materials and Methods

### DNA plasmid preparation and immunizations

Codon-optimized DNA plasmids (pVAX1 backbone) encoding toxin A RBD (residues 1848-2710, pRBDA) and toxin B RBD (residues 1851-2366, pRBDB) from *C. difficile* strain VPI10463 were constructed (GeneArt) as previously described(46). Briefly, putative N-linked glycosylation sites were removed through the substitution of glutamine for asparagine. Similarly, a codon-optimized DNA plasmid (pVAX1 backbone) encoding murine ADA was also constructed (Genscript) as previously described(39). Female and male young adult (6-8 weeks) and aged (72-76 weeks) C57BL/6J mice (Charles River) were used for vaccination and challenge studies. Aged C57BL/6J mice were obtained from the NIA Aged Rodent Colonies, which maintain barrier-raised, specific-pathogen-free mice specifically for aging research. These colonies provide access to well-characterized aged cohorts that support reproducible geroscience-focused studies. Aged animals were selected according to established geroscience guidelines(49). Our selection of 72–76-week-old mice align with established recommendations for modeling mid-to-late-life immune dysfunction while maintaining adequate health for infection and vaccination studies. Mice were immunized in the left tibialis anterior muscle with 5µg pRBDA + 5µg of pRBDB with or without 10µg pADA. Control mice were immunized with 20µg of the empty backbone vector, pVax, to match total DNA in vaccinated mice. Immediately after vaccine injection, *in vivo* electroporation was performed using a CELLECTRA^TM^ electroporator set to two pulses at a constant 0.2Amp current (Inovio Pharmaceuticals) according to previously reported tolerance guidelines(50, 51). Mice were immunized twice over the course of 28 days with the second immunization occurring at 28 days post the primary immunization. All mice were housed separated by sex and age in a temperature-controlled, light-cycled, specific-pathogen-free facility at Drexel University College of Medicine, no bedding was exchanged during the murine experiments. All murine work and protocols were approved by the appropriate Institutional Animal Care and Use Committee and Environmental Biosafety Committee.

### Mouse sacrifice, sample collection, and tissue harvest

At the indicated timepoints, mice were euthanized. For euthanization, mice were first given an 800mg/kg dose of avertin via intraperitoneal (IP) injection, and upon the lack of motor response to toe pinches, a terminal cardiac puncture was performed followed by cervical dislocation. Next, tissues, including spleen, and draining inguinal and popliteal lymph nodes (dLNs) were collected from each mouse and processed into single-cell suspensions (70uM strainer for splenocytes, 100uM strainer for lymph nodes) with complete RPMI media (10% FBS + 1x penicillin-streptomycin) for further use in cellular assays. Collected blood was placed in minicollect serum gel tubes (Grenier-Bio) and centrifuged at 16,000 rpm for 10 minutes at 4°C to separate out the serum. Serum was then frozen at -80°C until further use.

### ELISA assays

ELISAs were performed using flat, 96-well high-binding polystyrene plates (Corning Inc., Cat: 3590) to quantify *C. difficile* toxin A- and B-specific IgG in serum from vaccinated mice. Plates were first coated with 100uL of 0.5ug/mL of toxoid A (Enzo Life Sciences, Cat: BML-G145) or toxoid B (Enzo Life Sciences, Cat: BML-G155) overnight at 4°C. Plates were then blocked with blocking buffer (3% BSA, 0.1% tween-20, PBS) for 2 hours at room temperature. Mouse serum and standards (Clostridium difficile Toxin a Monoclonal Antibody A73H, Invitrogen, Cat: MA5-18192; Clostridium difficile Toxin B Monoclonal Antibody A13I, Invitrogen, Cat: MA1-7412) were then serially diluted in reagent diluent (1:100 of blocking buffer in PBS), added to the plate at 100uL/well, and incubated overnight at 4°C. Plates were washed (0.1% tween-20 in PBS) and incubated with 100uL/well goat anti-mouse HRP secondary antibody (1:10,000, Columbia Biosciences, Cat: HRP-112) for 1.5 hours at room temperature. Plates were washed and developed with TMB Ultra substrate (Thermo Fisher) for 75 seconds before adding stop solution (1M H_2_SO_4_). Plates were read on BioTek ELISA plate reader at 450nm.

### Activation-induced marker (AIM) and intracellular cytokine staining (ICS) assays

Splenocytes from vaccinated mice were plated in 96-well U-bottom plates at 1 x 10^6^ splenocytes per well and stimulated at 37°C for 6 or 18 hours with PMA, DMSO, or Toxin A and B RBD peptide megapools (1ug/mL/megapool) for ICS and AIM assays, respectively. Peptide pools for RBDA (residues 1848-2710) and RBDB (residues 1851-2366) were generated as separate pools of 15mers with 11 residue overlap (Genscript). For ICS, all stimulation conditions included Protein Transport Inhibitor Cocktail (eBioscience, Cat: 00-4980-93) and FITC-conjugated anti-CD107a (BioLegend, Clone:1D4B, Cat: 121606). Following stimulation, cells were transferred to 96-well V-bottom plates and stained with Live/Dead Aqua Fixable Viability Stain (Invitrogen, Cat: L34957) and anti-mouse CD16/32 (BioLegend, Clone: 93; Cat:101319) for 20mins at 4°C. For ICS surface staining, cells were incubated for 30 minutes at 4°C with the following antibodies: CD3-AF700 (BioLegend, Clone: 17A2, Dilution: 1:50, Cat:100216), CD4-APC-Cy7 (BioLegend, Clone: GK1.5, Dilution: 1:50, Cat:100414), CD8-BV711 (BioLegend, Clone: 53-6.7, Dilution: 1:30, Cat:100748), CD62L-PE/Dazzle594 (BioLegend, Clone: MEL-14, Dilution: 1:33, Cat:104448), CD44-PerCP/Cy5.5 (BioLegend, Clone: IM7, Dilution: 1:65, Cat:103032), PD1-BV421 (BioLegend, Clone: RMP1-30, Dilution: 1:50, Cat:109121), and CXCR5-BV785 (BioLegend, Clone: L138D7, Dilution: 1:20, Cat:145523). Following ICS surface staining, cells were washed and resuspended in 100µL Foxp3 / Transcription Factor Staining Buffer Set Perm/Fix solution (eBioscience, Cat:00-5523-00) for 1 hour at 4°C. Cells were intracellularly stained for cytokines with the following antibodies in 1x perm/wash buffer (eBioscience, Cat:00-5523-00) for 45 minutes at 4°C: IFNγ-APC (BioLegend, Clone: XMG1.2, Dilution: 1:100, Cat:505810), TNFα-BV650 (BioLegend, Clone: MP6-XT22, Dilution: 1:40, Cat:506333), IL-2-PE/Cy7 (BioLegend, Clone: JES6-5H4, Dilution: 1:100, Cat:503832), and BCL6-PE (BD, Clone: K112-91, Dilution: 1:20, Cat:561522).

For AIM surface staining, cells were incubated for 45 minutes at 4°C with the following antibodies: CD3-AF700 (BioLegend, Clone: 17A2, Dilution: 1:50, Cat:100216), CD4-APC-Cy7 (BioLegend, Clone: GK1.5, Dilution: 1:50, Cat:100414), CD25-PE/Cy5 (BioLegend, Clone: PC61, Dilution: 1:80, Cat:102010), OX40-BV421 (BioLegend, Clone: OX-86, Dilution: 1:100, Cat:119411), PD1-PE/Cy7 (BioLegend, Clone: RMP1-30, Dilution: 1:50, Cat:109110), PD-L1-BV605 (BioLegend, Clone: 10F.9G2, Dilution: 1:100, Cat:124321), CXCR5-APC (BioLegend, Clone: L138D7, Dilution: 1:20, Cat:145506), and 41BB-PE (BD, Clone: 1AH2, Dilution: 1:33, Cat: 558976).

All samples were then run on a BD LSR flow cytometer, and data were analyzed using FlowJo software v10.10.0 (BD Biosciences). The gating strategies for ICS and AIM data are provided in Supplementary Figure 1A, 3A.

### Switched/GC and Toxin-Specific B cell flow cytometry

Recombinant toxin A and toxin B probes were prepared and fluorescently-labelled with PE and AF647 as previously described(34). Single-cell suspensions of splenocytes from vaccinated mice were added to 96-well V-bottom plates at 2 x 10^6^ splenocytes per well. Cells were stained for viability in the presence of Live/Dead Aqua (Invitrogen, Dilution: 1:1000, Cat: L34957) and anti-mouse CD16/CD32 (BioLegend, Clone: 93, Dilution: 1:1000, Cat:101319) for 20mins at 4°C. Cells were then washed and stained for 1hr on ice in 100µL of antibody cocktail containing 1uL of fluorescently-labelled toxin A/B probes and the following conjugated anti-mouse antibodies: CD3-AF700 (BioLegend, Clone: 17A2, Dilution: 1:50, Cat:100216), CD19-BV605 (BioLegend, Clone: 6D5, Dilution: 1:50, Cat: 115530), IgD-PE/Dazzle (BioLegend, Clone: 11-26c.2a, Dilution: 1:50, Cat: 405742), IgM-BV786 (BD Horizon, Clone: R6-60.2, Dilution: 1:50, Cat: 564028), GL7-AF488 (Invitrogen, Clone: GL7, Dilution: 1:200, Cat: 53-5902-82), and CD38-BV421 (BioLegend, Clone: 90, Dilution: 1:100, Cat: 102732). All samples were then run on a BD LSR flow cytometer, and data were analyzed using FlowJo software v10.10.0 (BD Biosciences). The gating strategy is provided in Supplementary Figure 2A.

### TcdB_1961_ Tetramer labeling

Single cell suspensions from splenocytes and dLNs (popliteal and inguinal) were incubated with the TcdB_1961_ MHCII-I A^b^ tetramer(52) at 18 µg/mL in a 96 well round bottom plate for 2 hours at 37°C. Following tetramer labeling, cells were prepared for flow cytometry using a standard protocol. Briefly, cells were washed in PBS and stained with Zombie UV (BioLegend, Dilution: 1:500 Cat: 423107) or Live/Dead Aqua (Invitrogen, Dilution: 1:500, Cat: L34957) at room temperature for 10 minutes. Cells were washed with FACS buffer (1x PBS, 1% BSA, 1mM EDTA, 50µg/mL Sodium Azide) and blocked in blocking buffer (FACS Buffer with anti-16/32 [BD Biosciences] and rat IgG [Sigma]) for 10 minutes at 4C. Next, antibodies to surface antigens were added and incubated at 4C for 30 minutes. Antibodies included CD4-BUV395 (BD Biosciences, Clone: RM4-5, Dilution: 1:400, Cat: 568375), CD62L-BUV805 (BD Biosciences, Clone: MEL-14, Dilution:1:300, Cat: 569201), CD19-BV650 (BioLegend, Clone: 6D5, Dilution: 1:200, Cat: 115541), CD3-PerCP-Cy5.5 (eBioscience, Clone: 145-2C11, Dilution: 1:300, Cat: 35-0031-82), CD3-FITC (BioLegend, Clone: 500A2, Dilution: 1:200, Cat: 152304), CD5-PerCP-Cy5.5 (eBioscience, Clone: 53-7.3, Dilution: 1:300, Cat: 45-0051-82), CD5-FITC (BioLegend, Clone: 53-7.3, Dilution: 1:300, Cat: 100606), CD44-AF700 (BioLegend, Clone: IM7, Dilution: 1:200, Cat: 103026), CD44-PerCP-Cy5.5 (eBioscience, Clone: IM-7, Dilution: 1:500, Cat: 45-0441-82), PD-1-PE-Cy7 (eBioscience, Clone: J43, Dilution: 1:200, Cat: 25-9985-82), and CXCR4-APC (BioLegend, Clone: L276F12, Dilution: 1:50, Cat: 146508). For T_FH_ staining, cells were incubated with a biotinylated CXCR5 antibody (eBioscience, Clone: SPRCL5, Dilution: 1:50, Cat: 13-7185-82) for 45 minutes at 4C and washed before the addition of fluorophore conjugated streptavidin-BV421 (BioLegend, Cat: 410505) with all other surface antibodies. Following antibody staining, cells were washed twice in FACS buffer and fixed using 2% paraformaldehyde solution for 20 minutes at 4C. Cells were washed two more times in FACS buffer before FACS analysis. Cells were analyzed on a Symphony A3 lite (BD Biosciences), and data was analyzed using FlowJo Software. The gating strategy is provided in Supplementary Figure 1E, 4A.

### ELISpot assay

For detection of RBDA/B-specific IFNγ production, IFNγ ELISpot kits (R&D) were utilized and performed according to the manufacturer’s direction. Briefly, 96-well PVDF membrane (Mabtech) were coated with anti-mouse IFNγ capture antibodies overnight. Following washing with PBS, 250,000 cells/well were plated and stimulated with either DMSO as a negative control, RBDA/B (1µg/ml total final concentration), or Cell Activation Cocktail (BioLegend) as a positive control overnight. The following day, cells were removed, and plates were washed with PBS and incubated for 2 hours at room temperature with anti-mouse IFNγ secondary antibody. Streptavidin-ALP was then added, and plates were again incubated for 2 hours at room temperature. Finally, spots were developed using BCIP substrate for 15 minutes at room temperature, and spot-forming units (SFUs) were scanned and counted using a CTL ImmunoSpot S5 Core Analyzer (Cellular Technology Limited CTL).

### Toxin Neutralization Assay

To determine the neutralization capacity of vaccine-induced antibodies in mouse sera, Vero cells were propagated in complete DMEM (cDMEM) and plated at 8,000 cells/well in Nunc MicroWell 96-Well Black Optical-Bottom Plates (Thermo Scientific, Cat: 12-566-70). Following 37°C incubation for 24 hours (∼90% confluency), serum dilutions for individual mice were prepared and incubated with 60ng/mL Toxin A (List Biological Laboratories Inc., Cat: 152C) for 90 minutes at 37°C. As negative and positive controls, cDMEM and 100ug/mL Actoxumab (Invitrogen, Cat: MA5-41961) were incubated with 60ng/mL Toxin A, respectively. Supernatant was removed from the seeded Vero cells and 100uL of serum/Toxin A samples or controls were added to wells in duplicate. Individual plates also included control wells incubated with cDMEM alone for subsequent analysis of neutralization. After 72 hours of incubation at 37C, plates were imaged at a 4X Brightfield mode on a BioTek Cytation 5. To quantify neutralization, 100uL of CellTiterGlo2.0 reagent (Promega, Cat: G9241) were added to individual wells, mixed on an orbital shaker for 2 minutes (500 rpm), and incubated for an additional 10 minutes at room temperature for luminescence stabilization. Plates were read on a Cytation 5 plate reader to quantify luminescence. Neutralization was evaluated by calculating percent relative viability ((Luminescence RLU of sample/RLU cDMEM only control) *100).

### CD196 Spore Challenge and CFU Quantification

Prior to challenge, mice were pre-screened for *C. difficile* as previously described(53). Briefly, fresh fecal pellets were collected and resuspended in deoxygenated *C. difficile* selective broth to enrich for *C. difficile* for 24 hours. Ten-fold dilutions were plated on selective agar and incubated at 37C overnight to look for colony growth. Vaccinated young and aged mice were treated with antibiotic water containing vancomycin (0.33g/L), metronidazole (0.25g/L), and neomycin (0.33g/L) for 96 hours. Subsequently, mice were given regular water for 24 hours then injected intraperitoneally with clindamycin (250ug/mouse) on day 6. Mice were challenged the following day with 10^3^ spores of *C. difficile* strain CD196 spores diluted in 200uL sterile UltraPure water. Weight, morbidity, and mortality were monitored daily for 13 days(54, 55). Morbidity was scored based on coat appearance, posture, and mobility using a on a scale of 0, +1, and +2 representing normal, mild, and severe, respectively. Mice reaching >80% weight-loss or a morbidity score of +4 were euthanized prior to the experimental endpoint.

*C. difficile* burden (CFU/g) was quantified in fecal pellets from challenged mice on days 1 and 7 post-infection. Briefly, fecal pellets were collected and weighed for individual mice. In an anaerobic chamber, pellets were then homogenized in 1mL sterile PBS for subsequent 10-fold serial dilution preparation and plated on cc Brain Heart Infusion Supplemented (ccBHISTA) plates(55). Plates were incubated in the anaerobic chamber overnight at 37C, and colonies were counted the following day to enumerate CFU/g per mouse.

### Adenosine and metabolite measurements in dLNs

Popliteal and inguinal lymph nodes (LNs) were harvested from vaccinated mice following challenge (at time sacrifice or study endpoint) and placed in pre-weighed 1.5mL Eppendorf tubes. Tubes containing dLNs were immediately re-weighed and snap-frozen with 2-propanol on dry ice for at least 5 minutes before transfer to -80C for storage. Frozen lymph nodes were homogenized with bead-beating for 5 minutes in 100μL acetonitrile and water (1:1) and centrifuged. Supernatant was diluted 1:10 in acetonitrile, and adenosine was measured using liquid chromatography-mass spectrometry (LC/MS) on a Waters Acquity I-Class UPLC system coupled to a Synapt G2Si HDMS mass spectrometer in positive ion mode. LC separation was performed on a Merck SeQuant ZIC-cHILIC 3 µm 2.1x100mm column (maintained at 40 °C) using an 0.4 ml/min gradient of A/B 10/90 to 40/60 in 5 minutes, to 90/10 in 1 minute, followed by washing and reconditioning the column. Eluent A is 20 mM ammonium formate in water and B is acetonitrile. Conditions on the mass spectrometer were as follows: capillary voltage 0.5 kV, sampling cone 40, source offset 80, source 120 °C, desolvation 250 °C, cone gas 0, desolvation gas 1000 L/h, nebulizer 6.5 bar. Low energy data was collected between 100 and 1500 Da at 0.2 sec scan time. Using Waters Unifi 1.9, masses were extracted from the TOF MS TICs using an abs width of 0.05 Da, and peaks integrated. Sample readouts were normalized by weight (units/gram).

### Statistical analysis

Statistical analyses were performed using GraphPad Prism v9.5.1. Data distributions were assessed for normality, and parametric or nonparametric tests were applied accordingly. Comparisons involving more than two independent variables were analyzed using two-way ANOVA followed by Fisher’s least significant difference (LSD) post-hoc tests when the overall ANOVA was significant. Comparisons involving a single independent variable were performed using either Welch’s t test or the Mann–Whitney U test, as appropriate.

Survival differences were evaluated using Kaplan–Meier analysis with log-rank tests. Longitudinal weight-loss and morbidity scores were analyzed using mixed-effects models with Geisser–Greenhouse correction. All tests were two-sided, and a p value <0.05 was considered statistically significant. Data points represent individual animals unless otherwise indicated.

## Results

### C. difficile DNA vaccination design with or without pADA in young and aged mice

The immunogenicity of our *C. difficile* toxin A and B RBD DNA vaccine has not been evaluated in an aging model(46). Additionally, the effects of pADA co-immunization in an aged vaccination model has only been investigated in a virus-specific context with a particular focus on CD8+ T cells(40). Accordingly, we sought to investigate the immunogenicity of our DNA vaccine in aged mice and determine whether pADA co-immunization could restore age-associated defects in vaccine-induced immunity. Vaccine plasmids expressing toxin A RBD, toxin B RBD, and murine ADA are outlined in Figure 1A-B and were generated as previously described(39, 46). Young (6-8 weeks) and aged (72-76 weeks) mice were vaccinated twice at 0 and 28 days with 20ug pVax (control), 10ug pRBDA/B (referred to as pRBD hereafter), or 10ug pRBDA/B + 10ug pADA (referred to as pRBD/pADA hereafter). Spleens and serum were collected at ten days post-boost for subsequent cellular and humoral analyses (Figure 1C). This design enabled comparison of CD4 T cell and antibody responses between age groups and elucidation of ADA-dependent changes.

**Figure 1.**
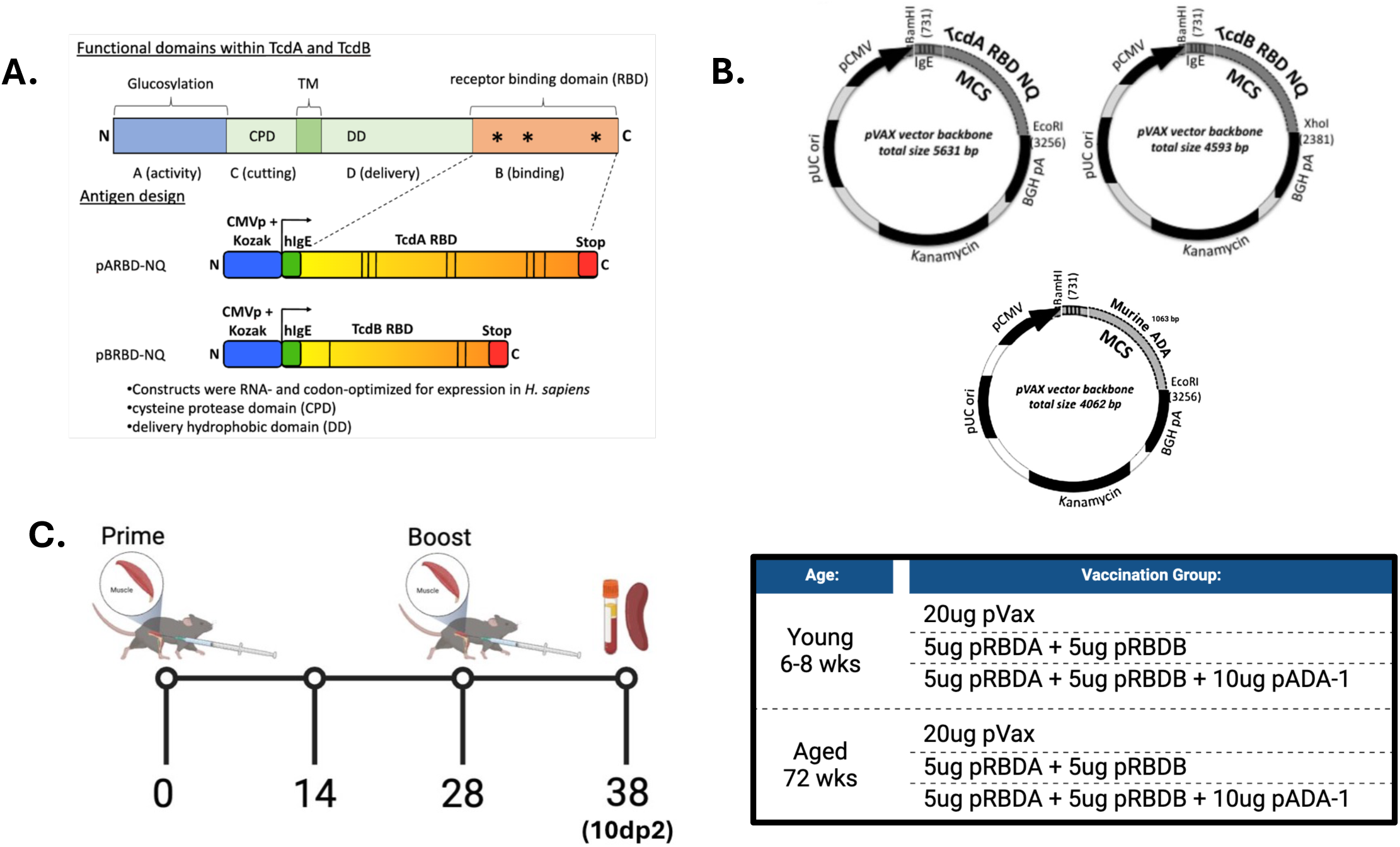
DNA vaccine design and immunization schedule. (A, B) Vaccine plasmid design (left) and constructs (right) encoding C. difficile toxin receptor binding domains or murine adenosine deaminase. (C) Experimental timeline where young (6-8 weeks) and aged (72-76 weeks) mice were vaccinated twice with pVax (20ug), pRBDA+pRBDB (5ug/plasmid), or pRBDA+pRBDB+pADA (5ug/RBD plasmid + 10ug pADA) over 28 days. Spleens and serum were collected at ten days post-boost for further analyses. Data from three independent experiments were pooled for statistical analysis. Experiments were performed using age-matched mice for both males and females.

### pADA enhances CD4^+^ T cell activation and polyfunctionality in aged mice

We have previously shown in the context of young murine HIV-1 and COVID-19 DNA immunizations, pADA co-immunization elicited a dose-sparing enhancement in CD4 and CD8 T cell responses and polyfunctionality(39, 40). To further evaluate the immunomodulatory effects of ADA on post-vaccination aged CD4^+^ T cells, in conjunction with known age-related defects in T cell cytokine production(14), we sought to evaluate IFNγ, IL-2, and TNFα production as key functional markers of Th1 and memory T cell responses, which are critical modulators of vaccine-induced protective immunity(56). First, we evaluated antigen-responsive IFNγ production from splenic T cells by ELISpot, a common method to assess vaccine and adjuvant efficacy(57–59).

Aged pRBD-vaccinated mice exhibited considerably lower toxin-specific IFNγ SFU compared to young mice (p=0.0282) (Figure 2A). Notably, pADA restored these responses in aged mice (p=0.0452) to a level comparable to young, vaccinated mice. To expand these results beyond total T cell responsiveness, we examined CD4^+^ T cell cytokine production. Following peptide stimulation of splenocytes, intracellular cytokine staining (IFNγ, IL-2, TNFα) in CD4^+^ T effector memory (T_EM_) (CD62L-CD44+) was quantified using flow cytometry. We observed significantly lower frequencies of IFNγ+ (p<0.0001) and TNFα+ (p<0.0001) in CD4^+^ T_EM_ from aged mice in comparison to young (Figure 2B, D). These frequencies were significantly increased in aged mice co-immunized with pADA (p=0.0296, p=0.0352) (Figure 2B, D). Similarly, pADA enhanced the frequency of IL-2+ CD4^+^ T_EM_ in aged mice (p=0.033) (Figure 2C). Concurrently, the number of CD4^+^ T_EM_ cells was similar between aged pRBD and pRBD/pADA vaccination groups (Supplementary Figure 1C).

**Figure 2.**
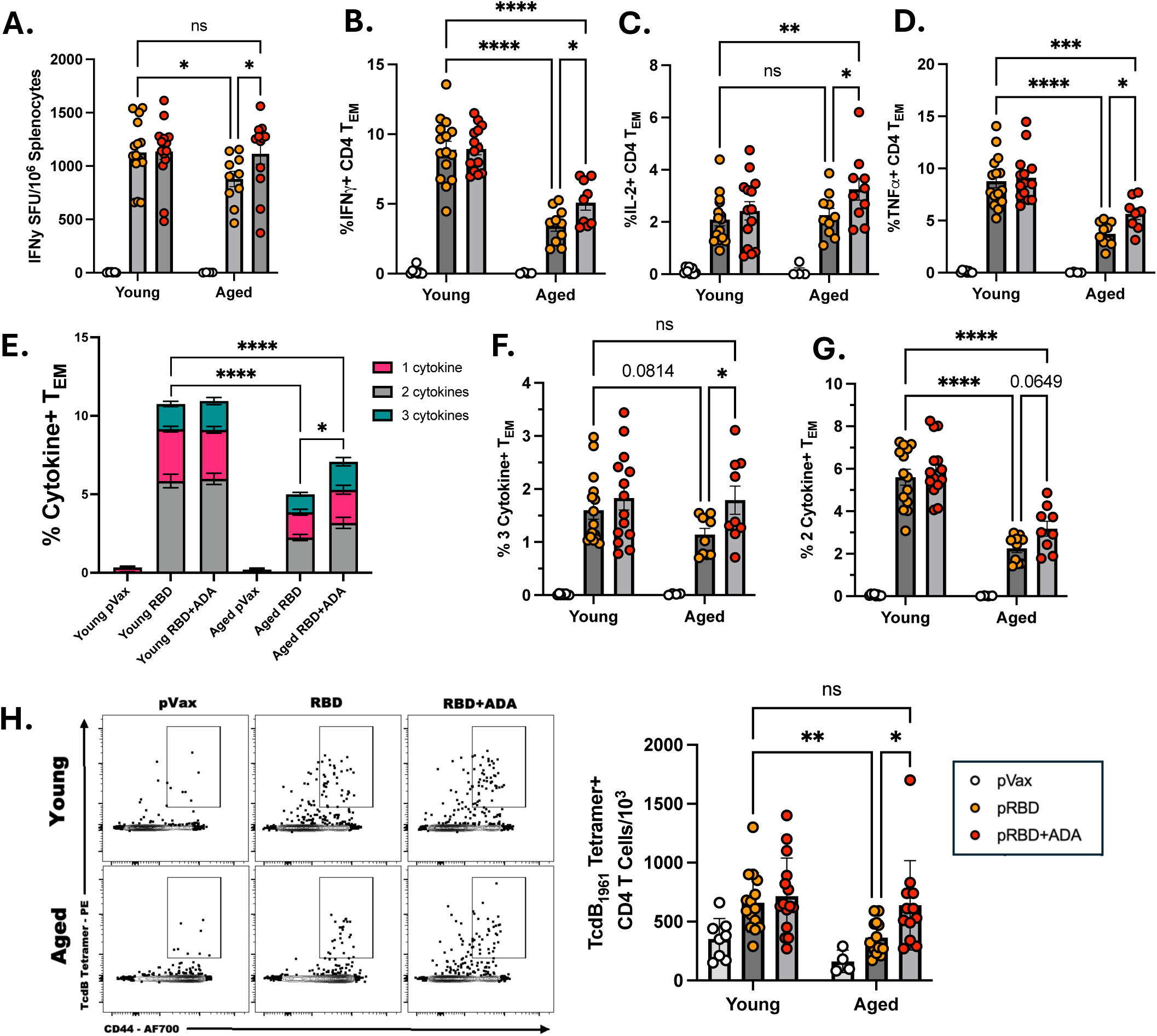
ADA restores RBD-specific CD4 T effector/memory cell responses in the aged mice. Young (6-8 weeks) and aged (72-76 weeks) mice were vaccinated with pVax (20ug), pRBDA+pRBDB (5ug/plasmid), or pRBDA+pRBDB+pADA (5ug/RBD plasmid + 10ug pADA) twice over 28 days. Spleens were harvested at ten days post-boost and processed for ELISpot and flow cytometry. (A) IFN**γ** production from toxin RBDA/B peptide megapool-stimulated splenocytes were measured by ELISpot and quantified as spot forming units (SFU) per 10^6^ cells. (B-D) Single and (E-G) polyfunctional IFN**γ**+, IL-2+, and TNF**α**+ T_EM_ (CD4+CD44+CD62L-) cells were quantified using flow cytometry following 6-hour RBDA/B peptide megapool stimulation. (H) TcdB_1961_ tetramer was used to quantify antigen-experienced (CD44+) CD4 T cells per 10^3^ splenocytes by flow cytometry. Data from three independent experiments were pooled for statistical analysis. Young pVax (n=8), Aged pVax (n=4), Young pRBD (n=15), Aged pRBD (n=10-13), Young pRBD+pADA (n=14-15), Aged pRBD+pADA (n=9-12). Error bars indicate mean and SEM. *P<0.05, **P<0.01, ***P<0.001 and ****P<0.0001 by 2-way ANOVA with Fisher’s LSD post-hoc test.

We further assessed polyfunctional cytokine responses (IFNγ, IL-2, TNFα) in peptide-stimulated CD4^+^ T_EM_ cells. Declining T cell polyfunctionality following vaccination is well-established in elderly humans and animal models(60–62). Consistent with these findings, pRBD-immunized aged mice displayed significantly lower cumulative polyfunctional responses (p<0.0001) and CD4^+^ T_EM_ expressing 2 cytokines (p<0.0001), with a trend towards lower IFNγ+IL-2+TNFα+ CD4^+^ T_EM_ (p=0.0814) (Figure 2E-G). Notably, pADA co-immunization significantly enhanced total polyfunctional responses (p=0.0332) (Figure 2E) and IFNγ+IL-2+TNFα+ CD4^+^ T_EM_ (p=0.0304) (Figure 2F), and augmented higher trending double-positive T_EM_ frequencies (p=0.0649) in aged mice compared pRBD alone (Figure 2G).

### pADA restores toxin-specific CD4^+^ T cell generation in aged mice

Given that dysregulated cytokine production is a hallmark of aging T cells, indirect measurement of vaccine responses by ICS can underestimate the total number antigen-specific T cells. Therefore, we next investigated whether pADA co-immunization improved the generation of toxin-specific CD4^+^ T cells in the spleen by direct detection. To do this, we quantified the number of vaccine-induced CD4^+^ T cells using a recently developed MHCII-I A^b^ tetramer specific for the immunodominant T cell epitope of the toxin B RBD domain (TcdB_1961_) (PMID: 41757099) (52), which is present in the coding region of the toxin B RBD plasmid. CD44+ was used as a marker to distinguish naïve CD4^+^ T cells from effector and memory (antigen-experienced) cells (Supplementary Figure 1E) (63–65). In pRBD-vaccinated groups, aged mice had significantly lower antigen-experienced TcdB_1961_-specific CD4^+^ T cells (p=0.0049) compared to young mice (Figure 2H), coinciding with the previously characterized age-dependent decline in antigen-specific CD4^+^ T cell production following vaccination(8, 66, 67). Importantly, pADA co-immunization significantly improved TcdB_1961_-specific antigen-experienced CD4^+^ T cell production in aged mice (p=0.0133) and restored the age-dependent reduction to a level comparable to young, vaccinated mice (Figure 2H). In agreement with the cytokine data, the tetramer data confirm pADA co-immunization expands vaccine-induced toxin-specific CD4^+^ T cells in aged mice to levels comparable to young.

### Antibody-mediated toxin A neutralization is restored by pADA

CD4^+^ T cells are necessary for robust, protective, and long-lasting antibody responses through cytokine production and direct contact with B cells. After observing enhanced vaccine-specific CD4^+^ T_EM_ cell responses, we quantified αTcdA and αTcdB IgG responses to determine if ADA improved total antibody production ten days post-boost. Only αTcdA IgG responses were significantly impaired in aged pRBD mice compared to young (p=0.0288), with similar titers between young and aged αTcdB IgG (Figure 3A-B). pADA co-immunized aged mice did not significantly differ from young pRBD and displayed a moderate 1.6-fold increase in median αTcdA IgG compared to pRBD-immunized aged mice (Figure 3A). No significant differences were observed in αTcdB titers, but median titers in young and aged pRBD/pADA groups exhibited 1.7-fold and 2.2-fold increases, respectively, compared to pRBD-vaccinated mice (Figure 3B).

**Figure 3.**
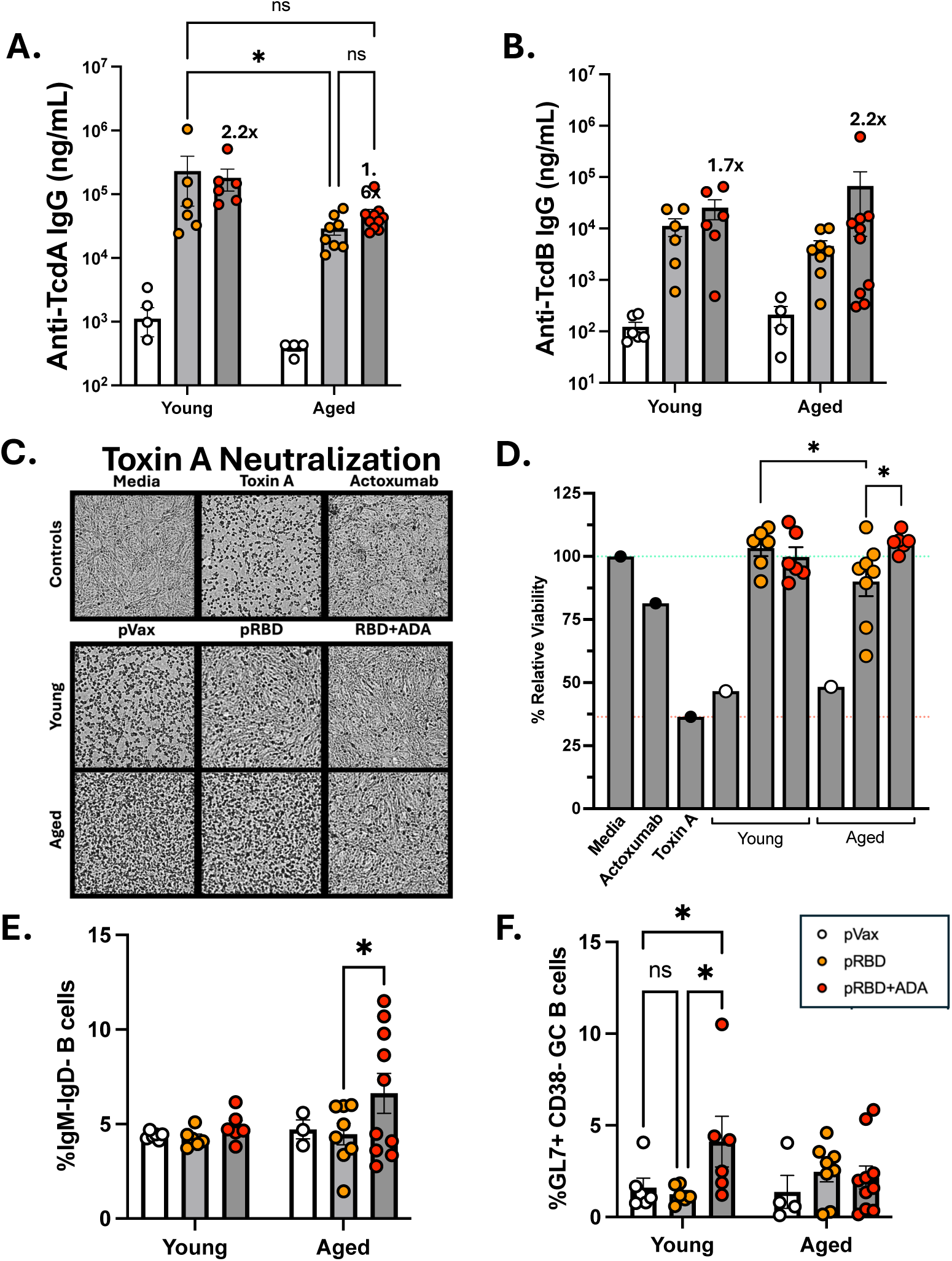
ADA restores the quality of humoral responses following booster immunization in aged mice. Serum and splenocytes were collected from twice-vaccinated mice at ten days post-boost. Quantification of (A) Toxin A- and (B) B-specific serum IgG by ELISA. Fold-change was quantified using median IgG. (C) Representative brightfield microscopy images (4X) and (D) percent relative viability of Vero cells quantified using CellTiterGlo luminescence readouts following incubation with toxin A and serum diluted 1:30 in PBS. Percent relative viability =(RLU Sample/RLU Media)x100. Frequency of (E) switched (IgM-IgD-) and (F) germinal center (GL7+CD38-) B cells in splenocytes quantified with flow cytometry. Data from three independent experiments were pooled for statistical analysis. (A, B, D) Values represent the mean of duplicate measurements for each animal. Young pVax (n=1-6), Aged pVax (n=1-4), Young pRBD (n=6-8), Aged pRBD (n=8), Young pRBD+pADA (n=6), Aged pRBD+pADA (n=6-10). Error bars indicate mean and SEM. *P<0.05, **P<0.01, ***P<0.001, and ****P<0.0001 by 2-way ANOVA with Fisher’s LSD post-hoc test.

Although antibody quantity represents an important correlate of protection for vaccine responses and *C. difficile* susceptibility, antibodies must also be functional to elicit protection. Antibody neutralization of toxins A and B are key functional correlates of protection against *C. difficile*-associated disease and recurrence by inhibiting toxin binding, internalization, and subsequent intoxication of cells(68–71). Moreover, a reduction in vaccine-elicited neutralizing antibodies represents another hallmark of aging immunosenescence(5, 39). Therefore, we next assessed if ADA altered the quality of antibodies produced by toxin-specific B cells by examining toxin A neutralization from the sera of immunized mice. Consistent with prior reports of immunosenescent vaccine responses, serum toxin A neutralization—measured as percent relative viability of seeded Vero cells—was significantly reduced in aged pRBD-vaccinated mice compared to young counterparts (p=0.0372) (Figure 3C, D). Importantly, pADA co-immunization significantly improved toxin A neutralization in aged mice (p=0.0162), restoring neutralization to a level comparable to young pRBD-vaccinated mice. As B cells are the ultimate source of antibody production, we examined if pADA had any effect on the B cell compartment. pADA significantly enhanced the frequency of class-switched B cells in aged mice (p=0.0249) (Figure 3F), but had no effect on the frequencies of GC B cells and toxin-specific switched B cells which were similar between aged vaccination groups (Figure 3E and Supplementary Figure 2B, C), suggesting the enhanced neutralization was not a result of B cell modulation. Collectively, these findings demonstrate that pADA co-immunization significantly enhances the functional quality of vaccine-induced antibodies, rather than quantity, in aged mice. In the absence of detectable changes within the B cell compartment, these data suggest that improved toxin A neutralization is likely mediated through ADA-dependent modulation of T_FH_ cells.

### pADA reverses impaired toxin-specific T_FH_ activation in aged mice

The induction of neutralizing antibody responses is a hallmark of vaccine responses, particularly in those targeting bacterial toxins. T_FH_ cells are critical mediators of vaccine-induced neutralizing antibody production by inducing B cell somatic hypermutation and class-switching(72, 73). Considering the previously described role of ADA in T_FH_ delineation and functionality(41), we sought to investigate whether pADA co-immunization impacted age-related T_FH_ cell dysfunction. First, we evaluated T_FH_ activation as an indicator of function at an earlier timepoint post-boost during peak T_FH_ responses (Figure 4A). Using a flow cytometry-based activation-induced marker (AIM) assay with an 18 hour peptide stimulation for T_FH_ enrichment(74), activation of vaccine-induced T_FH_ cells were evaluated by PD-L1, CD25, and OX40 co-expression(74, 75). Expression of these markers have been reported as functional indicators of antigen-induced GC T_FH_(75–77). Interestingly, the frequency of PDL1^+^CD25^+^ and CD25^+^OX40^+^ AIM+ toxin-specific T_FH_ cells were significantly lower in aged mice compared to young mice (p=0.0185 and p<0.0001, respectively) (Figure 4B, C). Importantly, pADA co-immunization significantly enhanced the frequency of PD-L1^+^CD25^+^ (p=0.003) and CD25^+^OX40^+^ (p=0.0398) co-expressing T_FH_ (Figure 4B, C). A similar trend was observed for single activation markers (PD-L1+, CD25+, OX40+) (Supplementary Figure 3B). Significantly higher PD-L1 expression was also observed in aged mice co-immunized with pADA (p=0.0005) (Supplementary Figure 3D). Taken together, the pADA-dependent restoration of antibody-mediated toxin-A neutralization in aged mice likely results from direct enhancement of vaccine-induced T_FH_ cell responsiveness and function.

**Figure 4.**
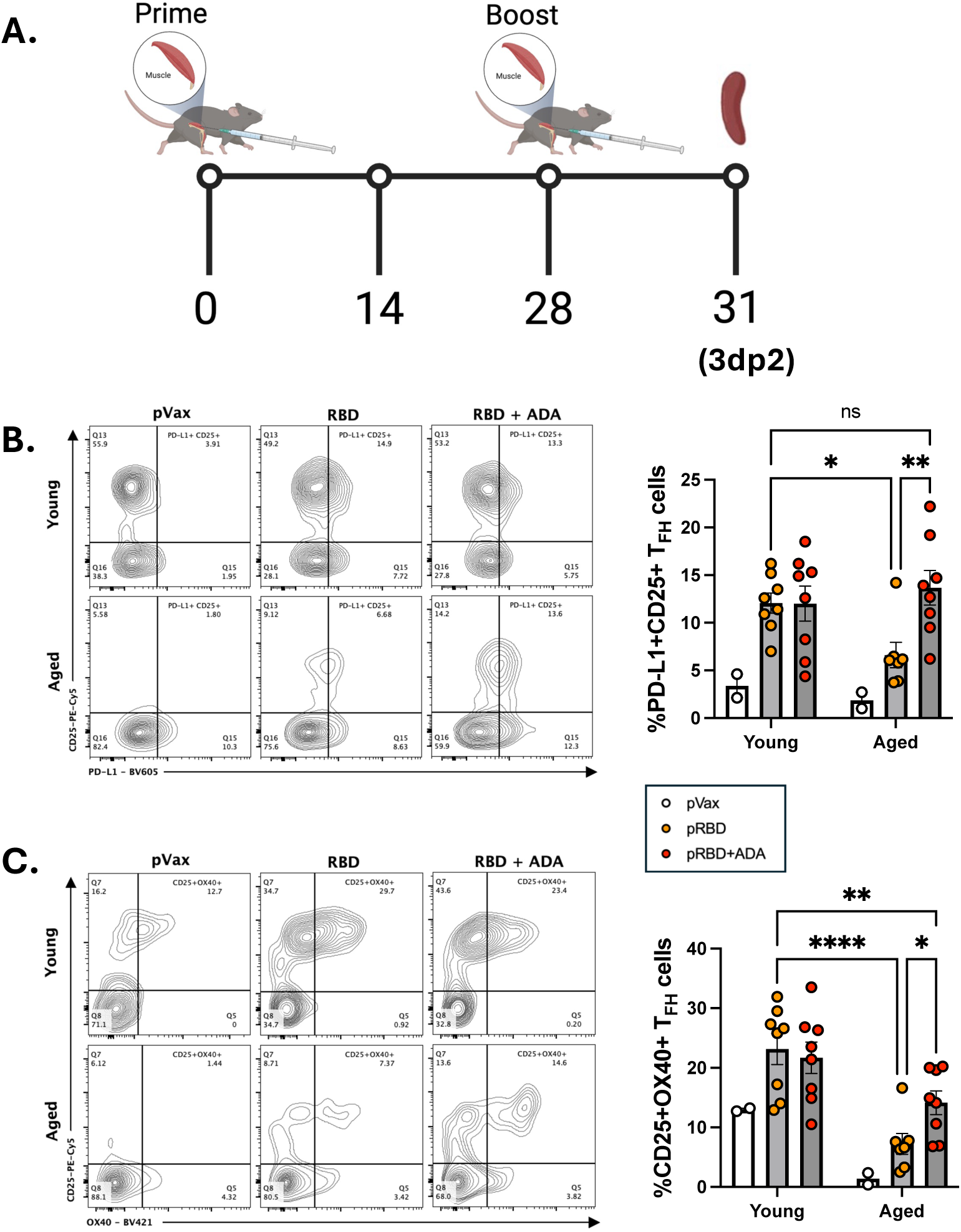
ADA restores antigen-specific T_TH_ cell activation in aged mice. (A) Experimental timeline where young (6-8 weeks) and aged (72-76 weeks) mice were vaccinated twice with pVax (20ug), pRBDA+pRBDB (5ug/plasmid), or pRBDA+pRBDB+pADA (5ug/RBD plasmid + 10ug pADA) over 28 days. Spleens were harvested and processed for flow cytometry at three days post-boost. (B) PD-L1+CD25+ and (C) CD25+OX40+ AIM-positive T_FH_ frequencies in splenocytes were measured using flow cytometry. Data from two independent experiments were pooled for statistical analysis. Experiments were performed using age-matched mice for both males and females. Young pVax (n=2), Aged pVax (n=2), Young pRBD (n=8), Aged pRBD (n=7), Young pRBD+pADA (n=8), Aged pRBD+pADA (n=8). (B, C) Error bars indicate mean and SEM. *P<0.05, **P<0.01, ***P<0.001 and ****P<0.0001 by 2-way ANOVA with Fisher’s LSD post-hoc test.

### pADA co-immunization reduces CXCR4+ GC T_FH_ cells in aged lymph nodes

In addition to active GC formation, vaccine-induced neutralizing antibody production requires GC T_FH_ light zone positioning for antigen-specific T_FH_-to-B cell interactions. The recent characterization of post-vaccination GC responses revealed that the expansion of CXCR4+ GC T_FH_ cells largely mediates age-associated decline in antibody quality(10). Mechanistically, age-related CXCR4 upregulation promotes T_FH_ mislocalization to the dark zone of GCs where functional B cell help is disrupted. Given the observed pADA-mediated restoration of antibody neutralization in aged mice, we postulated pADA co-immunization may reduce CXCR4 expression on GC T_FH_, thus promoting efficient GC vaccination responses. Based on the direct association between CXCR4 expression and GC T_FH_ distribution, we evaluated total and TcdB_1961_-specific GC T_FH_ in the draining popliteal and inguinal lymph nodes (dLNs) from immunized mice (Figure 4A). The total number of GC T_FH_ was unaltered by ADA within each age group, though aged mice exhibited higher overall GC T_FH_ cells similar to previous reports (Figure 5A)(40, 78). Comparable to the findings in Silva-Cayetano et al., higher total (p<0.0001) (Figure 5B) and TcdB_1961_-specific (p<0.0001) (Figure 5C and Supplementary Figure 4B) CXCR4+ T_FH_ were detected in aged mice vaccinated with pRBD. Importantly, pADA co-immunization reduced the number of both TcdB_1961_-specific (p=0.006) and total (p=0.0508) CXCR4+ GC T_FH_ populations in aged mice (Figure 5B-C). Notably, pADA co-immunization resulted in a two-fold reduction in CXCR4 surface expression on TcdB_1961_-specific GC T_FH_ cells in aged mice (p=0.0401) (Figure 5D), revealing a previously unrecognized role for ADA in regulating T cell CXCR4 expression. Comparable levels of non-specific (Figure 5A) and TcdB_1961_-specific GC T_FH_ (Supplementary Figure 4C-D) within age groups suggest ADA exerts expression-specific modulation of CXCR4, potentially through its enzymatic activity (Figure 5E).

**Figure 5.**
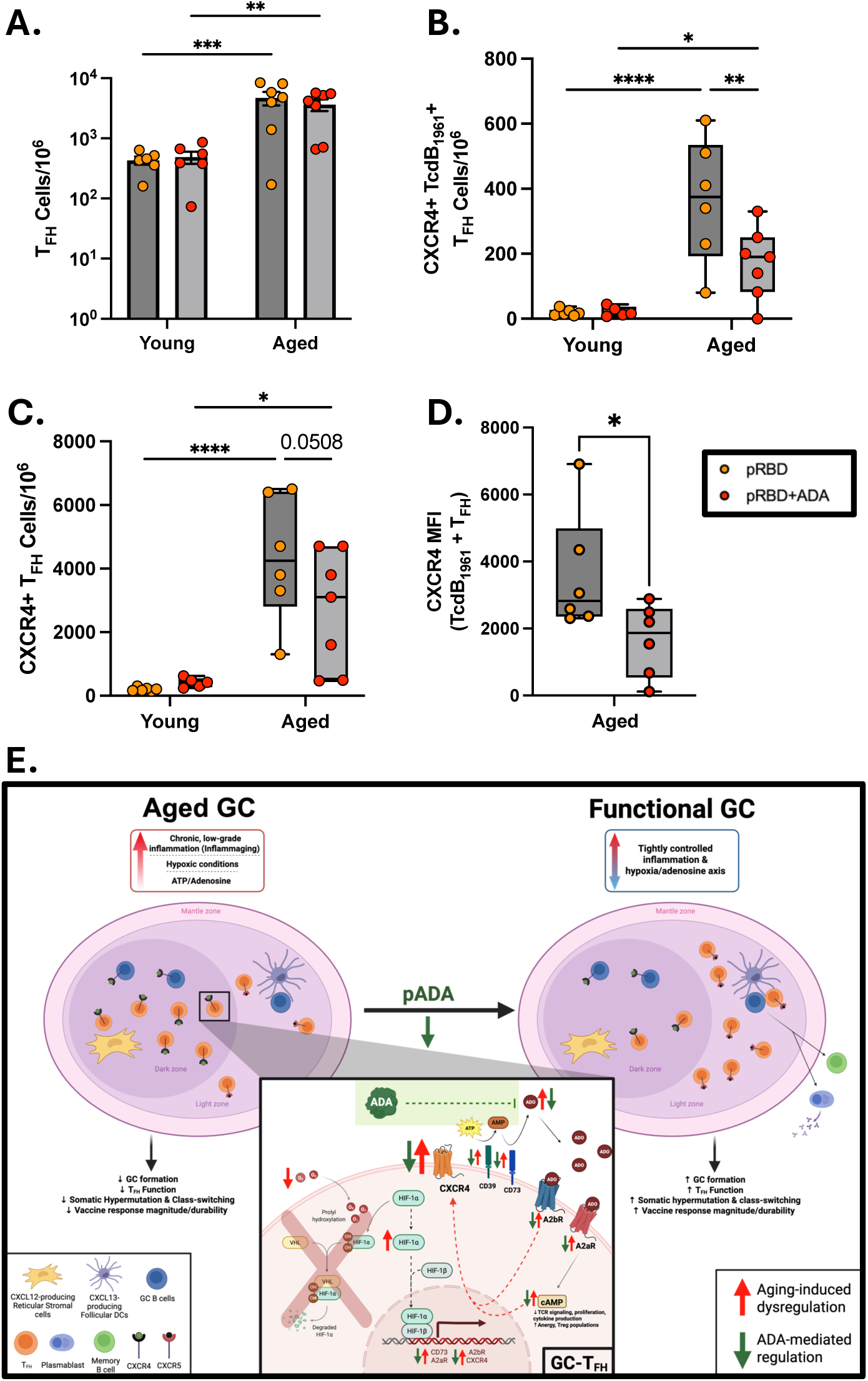
ADA reduces CXCR4 expression on aged antigen-specific GC Tfh isolated from popliteal and inguinal lymph nodes post-boost. Young (6-8 weeks) and aged (72-76 weeks) mice were vaccinated twice with pVax (20ug), pRBDA+pRBDB (5ug/plasmid), or pRBDA+pRBDB+pADA (5ug/RBD plasmid + 10ug pADA) over 28 days. Popliteal and inguinal lymph nodes were processed and analyzed using flow cytometry. (A) Total GC T_FH_ cells per 10^6^ lymphocytes, (B) CXCR4+ GC T_FH_ cells per 10^6^ lymphocytes, (C) CXCR4+ TcdB_1961_-specific GC T_FH_ cells per 10^6^ lymphocytes, and (D) CXCR4 mean fluorescence intensity on TcdB_1961_+ GC T_FH_ from aged mice were quantified. (E) Proposed model: ADA re-establishes metabolic regulation of adenosine in the aged GC microenvironment, dampening A2aR-cAMP and A2bR signaling in GC TFH, and lowering CXCR4 and improving T_FH_ localization/function and antibody quality in aged GCs. T_FH_, T follicular helper cells; FDC, follicular dendritic cells; CXCR4, C-X-C chemokine receptor 4; CXCR5, C-X-C chemokine receptor 5; A2AR, adenosine A2A receptor; cAMP, cyclic adenosine monophosphate; HIF-1α, hypoxia-inducible factor 1 alpha; CD39/CD73, ectonucleotidases; ↑, increased expression or production; ↓, decreased expression or production. Created with BioRender. (A) Error bars indicate mean and SEM and (B-D) minimum to maximum. Data from two independent experiments were pooled for statistical analysis. Young pRBD (n=1-6), Aged pRBD (n=6-7), Young pRBD+pADA (n=5-6), Aged pRBD+pADA (n=7). *P<0.05, **P<0.01, ***P<0.001 and ****P<0.0001 by (A-C) 2-way ANOVA with Fisher’s LSD post-hoc test and (D) Mann-Whitney U test.

### pADA co-immunization improves morbidity and mortality in aged mice following CD196 challenge

To determine if pADA-enhanced vaccine responses augment protection from infection, young and aged mice were challenged with 10^3^ CFU *C. difficile* CD196 spores 6 weeks post-boost as previously described (Figure 6A)(34). As expected, aged controls (pVax) exhibited greater susceptibility to infection (33.3% survival) compared to young mice (66.7% survival) (Figure 6C). Whereas young mice were completely protected by pRBD vaccination regardless of ADA, pRBD-immunized aged mice exhibited significantly lower survival compared to young mice (p=0.028), consistent with the known age-dependent decline in vaccine efficacy. Importantly, pADA co-immunization moderately improved overall survival in aged mice to a level statistically comparable to young mice. Beyond survival, vaccine efficacy can also be evaluated by disease burden (i.e. morbidity and weight-loss). Aged mice co-immunized with pADA displayed significantly lower weight loss (Figure 6D) and morbidity (Figure 6E) during early and peak infection (1-5dpi) compared to aged pRBD-immunized mice(48). Concurrently, aged mice co-immunized with pADA resembled young RBD-vaccinated mice in weight-loss and morbidity at all timepoints post-challenge (Supplementary Figure 5C, F), whereas aged mice vaccinated with RBD alone exhibited greater weight loss and morbidity (Supplementary Figure 5B, E). Examination of fecal pellets early (1dpi) and late (7dpi) showed ADA-specific improvements in aged mice were independent of *C. difficile* burden, as the control and vaccinated groups exhibited comparable CFU/g (Figure 6B). Young mice co-immunized with pADA curiously exhibited significantly lower CFU/g compared to control mice despite this vaccination model lacking spore-and vegetative cell-specific antigens. This data is the first to indicate pADA co-immunization can significantly reduce disease severity in aged, vaccinated mice.

**Figure 6.**
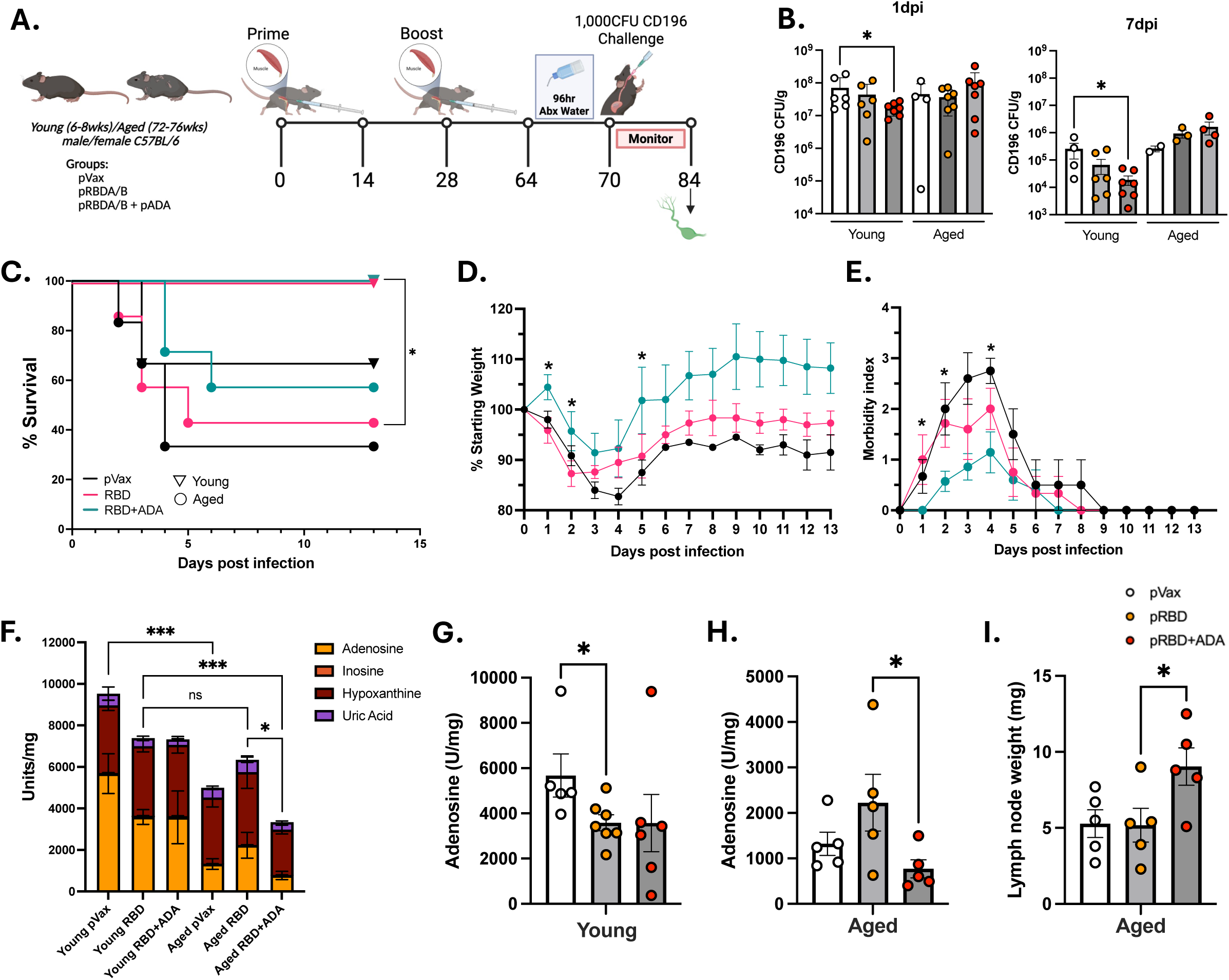
Mortality, weight loss, and morbidity in vaccinated young and aged mice following CD196 challenge. (A) Young (6-8 weeks) and aged (72-76 weeks) mice were vaccinated twice with pVax (20ug), pRBDA+pRBDB (5ug/plasmid), or pRBDA+pRBDB+pADA (5ug/RBD plasmid + 10ug pADA) over 28 days, rested for 6 weeks, then challenged with 10^3^ CD196 spores by oral gavage. (B) *C. difficile* CD196 burden (CFU/g) was quantified in fecal pellets at days 1 and 7 post-infection. (C) Mortality, (D) weight loss, and (E) morbidity were tracked daily for 13 days post-challenge. (D, E) Weight-loss and morbidity data depicts aged groups only. Morbidity scores were assigned based on established clinical scoring criteria (see Methods). (F-H) Adenosine, inosine, hypoxanthine, and uric acid were measured in popliteal and inguinal lymph nodes post-challenge and normalized by weight. (I) Lymph node weights (mg) in vaccinated aged mice. (B, D-G) Error bars indicate mean and SEM. Experiments were performed using age-matched mice for both males and females. Young pVax (n=5-6), Aged pVax (n=5-6), Young pRBD (n=7), Aged pRBD (n=5-7), Young pRBD+pADA (n=6-7), Aged pRBD+pADA (n=7). (B, F-I) Error bars indicate mean and SEM. *P<0.05, **P<0.01, ***P<0.001 and ****P<0.0001 by (B, G, H) Mann-Whitney U test; (C) Kaplan-Meier curves with Log-Rank Test (survival); (D-E) Mixed-Effect Analysis with Geisser-Greenhouse Correction (Weight-loss, Morbidity Index); (F) 2-way ANOVA with Fisher’s LSD post-hoc test; (I) Welch’s t-test.

### pADA co-immunization corrects dysregulated adenosine regulation in aged draining lymph nodes

In contrast to human ADA, murine ADA is unable to bind CD26 for T cell co-stimulation(79). Based on ADA’s established enzymatic role in immunosuppressive adenosine catabolism, we hypothesized that pADA co-immunization restores functional T cell and humoral responses in aged mice by correcting age-associated defects in adenosine regulation within vaccine-draining lymph nodes. To evaluate this mechanism, we quantified adenosine and related purine metabolites in popliteal and inguinal draining lymph nodes (dLNs) from vaccinated aged mice following *C. difficile* challenge using LC/MS. In pRBD vaccinated young mice, antigen re-exposure was associated with a significant reduction in dLN adenosine relative to primary exposure (pVax) (p=0.0303), reflecting physiologic metabolic remodeling during productive recall responses (Figure 6F, G). In contrast, aged mice failed to downregulate adenosine following antigen re-exposure, instead exhibiting aberrant accumulation within dLNs.

Consistent with our hypothesis, pADA co-immunization significantly reduced adenosine levels in the dLNs of aged mice compared with pRBD alone (p = 0.0317; Figure 6H). This reduction occurred in the absence of a global depletion of downstream purine metabolites, including inosine, hypoxanthine, and uric acid, indicating that pADA selectively corrected excessive adenosine accumulation (Figure 6F, H). Notably, reduced adenosine levels in pADA co-immunized aged mice were accompanied by significantly increased dLN weights (p = 0.0484; Figure 6I), consistent with enhanced lymphoid cellularity and active germinal center responses. Comparison of pVax mice from both cohorts revealed higher total metabolite levels in young mice (p=0.0006; Figure 6F), suggesting effective immune recall is associated with dynamic regulation of adenosine rather than low absolute levels. These data indicate pADA co-immunization restored dysregulated adenosine production in aged mice, yielding adenosine dynamics more closely aligned with those observed in young animals.

## Discussion

The continuous, rapid expansion of the aging population worldwide has created a growing interest in the prevention and treatment of age-associated diseases. Therapeutic approaches aimed at regulating and restoring immune function are key in the ongoing efforts to extend human health and longevity. Therefore, in addition to unraveling the underlying mechanisms of immunosenescence, new approaches for immune modulation are necessary.

Prior work outlining the association of ADA with T_FH_ cell function, GC-T_FH_-B cell interactions, and neutralizing antibody production(39–42) during vaccination and chronic infection suggests a promising role of ADA-mediated restoration of aged immunity. More importantly, the imbalance of ADA/adenosine axis in aging has prompted us to investigate whether co-immunization with pADA could help reverse immunosenescence. In fact, apart from enhancing vaccine response quality and quantity in multiple models(39, 40, 42), ADA has emerged as a promising therapeutic target for either supplementation or inhibition across multiple contexts including leukemia chemotherapy, CAR T cell optimization and exhaustion resistance, and cardiovascular disease therapies(80–83). The dynamic influence of ADA across varying tissues and disease states highlights the importance of appropriate ADA regulation. With aging representing a driving factor of immunosenescent responses in many disease states and therapy failures, we sought to understand the modulatory effects of pADA co-immunization on age-dependent vaccine responses. Unlike the results described in a COVID-19 vaccination model in aged mice(40), this work aimed to understand the effects of ADA on immunosenescence on antigen-specific CD4^+^ T cell populations using a vaccine targeting bacterial antigens. To date, direct detection of *C. difficile* toxin–specific CD4^+^ T cells has been limited by the absence of peptide–MHC class II tetramers. Here, we are the first to utilize a recently developed TcdB-specific tetramer (TcdB_1961_)(52) to directly track antigen-specific CD4^+^ T cells in aged mice following immunization.

*C. difficile* infection represents a relevant model of significant age-associated susceptibility yet lacks clinically available vaccines. Importantly, employing a previously described model of murine DNA vaccination and *C. difficile* infection, this study recapitulated impaired immune function and challenge outcome associated with aging. Of note, aged mice exhibited significantly diminished production of TcdB_1961_-specific CD4^+^ T cells (Figure 2H) in addition to activation-induced cytokine production and polyfunctionality in CD4^+^ T_EM_ cells (Figure 2A-G) at ten days post-boost. At the same timepoint, humoral responses including anti-toxin A IgG (Figure 3A) and serum toxin A neutralization (Figure 3C, D) displayed significant age-associated decline. These findings coincide with previous reports of age-associated decline in vaccine immunity(84). Furthermore, earlier post-boost, toxin-specific T_FH_ from aged mice demonstrated significantly reduced frequencies of AIM+ cells (Figure 4B-C) and reduced expression of activation markers PD-L1 (Figure 4D), CD25, and OX40 (Supplementary Figure 3C, D). Finally, aged pRBD-vaccinated mice had significantly lower survival against *C. difficile* challenge compared to young mice, which exhibited complete protection (Figure 6C). These results collectively highlight the impact of immunosenescence on vaccine-elicited protection, thus enabling relevant investigation of ADA-mediated restoration of aged immune responses using young mice as physiological benchmark for maximal vaccine responsiveness.

Through this work, we have demonstrated a novel role of pADA co-immunization in restoring antigen-specific CD4^+^ T cell generation (Figure 2H) and T_FH_ activation, indicated by PD-L1, CD25, and OX40 expression (Figure 4B-C), in aged mice. Prior investigation assessing CD25 and OX40 AIM markers on antigen-specific T_FH_ from immunized mice corroborates these findings as indicative of functional improvement(74). Activation-dependent OX40 expression holds substantial relevance as a promoter of T_FH_ differentiation, survival, and helper function for B cells(85). Transient CD25 upregulation on reactivated GC T_FH_ has also been implicated in B cell helper function and immune memory development(86). Additionally, this data is the first to indicate ADA-mediated restoration of aged T_FH_ activation coincides with functional restoration of antibody-mediated toxin neutralization (Figure 3C, D). These findings are particularly striking as defective antigen-specific T_FH_ activation of B cells(9) and reduced GC T_FH_ formation(87) in aging are key mechanisms underlying GC dysfunction and neutralizing antibody generation(88, 89). Additional improvements were observed in aged mice receiving pADA co-immunization. For instance, ADA significantly enhanced toxin-specific T cell responses measured by ELISpot and ICS (Figure 2A-D). Polyfunctional CD4^+^ T_EM_ responses were also improved, although they did not fully reach the levels observed in young mice (Figure 2E-G). Notably, the number of CD4^+^ T_EM_ cells were comparable among vaccinated mice of the same age group (Supplementary Figure 1C), suggesting ADA primarily augments antigen-specific functional capacity rather than broadly expanding the effector memory compartment. Decisively, expansion of antigen-specific CD4^+^ T cells with polyfunctional cytokine secretion are a hallmark of vaccine-induced T cells capable of supporting durable cellular memory and B cell proliferation(5, 7, 90). Thus, our findings in two major determinants of vaccine failure in older populations – functional CD4^+^ T cell generation and T_FH_ activation – underscore ADA’s ability to enhance protective immunity even at advanced age.

Furthermore, serum anti-toxin A and B IgG production showed a slight, but non-significant increase with pADA co-immunization (Figure 3A-B), suggesting ADA primarily biases toward enhanced antibody functionality rather than quantity in aged mice. Elevated frequencies of switched B cells, but not GC B cells, were detected in aged pADA co-immunized mice (Figure 3E, F). However, toxin-specific switched B cells were unchanged (Supplementary Figure 2B, C), indicating ADA primarily exerts its immunomodulatory function on the T cell compartment. Collectively, these data suggest pADA reverses immunosenescence in antibody quality by facilitating antigen-specific GC T_FH_-mediated B cell help, including somatic hypermutation, enabling production of functional, neutralizing antibodies. When translated to an infection model, pADA co-immunization resulted in a near 15% increase in survival compared to pRBD-vaccinated aged mice (Figure 6C). Strikingly, aged mice co-immunized with pADA exhibited weight-loss and morbidity analogous to young pRBD-vaccinated mice (Supplementary Figure 5C, F), whereas aged pRBD-vaccinated mice were significantly less protected (Supplementary Figure 5B, E). Of the pADA co-immunized aged mice that succumbed to infection, delayed mortality extending out to day 6 was observed (Figure 6C). Additionally, significantly lower weight loss and morbidity scores were observed on days 1, 2, 4, and 5 post-infections in pRBD/pADA-immunized aged mice (Figure 6D-E), indicating pADA reduced disease severity at early and recovery timepoints post-infection. These findings indicate that ADA-mediated correction of age-associated immune defects conferred measurable protection following challenge, directly connecting immune restoration to improved clinical outcome.

To identify a potential mechanism of ADA-mediated restoration of GC T_FH_ and neutralizing antibody responses, we investigated a recently described driver of immunosenescent vaccine responses, CXCR4. Specifically, age-associated upregulation of CXCR4 on GC T_FH_ cells was shown to impair functional T_FH_-B cell interactions by promoting CXCL12-induced migration to the dark zone(10). Spatial mislocalization prevents T_FH_ from providing B cell help in the GC light zone via induction of isotype switching and somatic hypermutation, thus impairing neutralizing antibody generation. Because pADA restored aged-related impairments in antibody-mediated toxin neutralization and T_FH_ activation, we postulated pADA restores CXCR4-induced spatial mislocalization of GC T_FH_. Indeed, pADA significantly reduced the age-dependent increase in CXCR4+ GC T_FH_ cells and CXCR4 surface expression in the dLNs of vaccinated aged mice (Figure 5B-D). Combined, these findings suggest pADA co-immunization may prompt enhanced GC T_FH_ function by altering CXCR4 expression. Importantly, while aged mice had higher total and toxin-specific GC T_FH_ cells compared to young mice, the number of cells within each age group was similar regardless of vaccination group (Figure 5A and Supplementary Figure 4D). This data supports expression-specific modulation of CXCR4 by pADA co-immunization rather than alteration of T_FH_ cell quantity during aged vaccine responses. Further investigation is required to determine the mechanism by which ADA can modulate CXCR4 expression in T_FH_.

One possible explanation involves ADA’s primary role in adenosine metabolism. In immunocompetent individuals, vaccination induces the rapid expansion of germinal center lymphocytes, creating a hypoxic environment that promotes high adenosine production to quell excessive inflammation. However, dysregulation of this hypoxia-adenosine axis can drive potent immunosuppression, best exemplified by hypoxia-induced CD39-dependent T cell exhaustion in tumor microenvironments (TME)(91). Analogous to the TME, aging drives the accumulation of terminally-differentiated effector T cells(92, 93) through elevated expression of adenosine-producing CD39 and the adenosine receptor A2aR(44), which are known inhibitors of T_FH_ differentiation and function(94). Consequently, aged T cells demonstrate higher adenosine production (43) and a CD39+ apoptosis-prone phenotype when activated(44). Moreover, adenosine signaling through A2aR and A2bR has been shown to inhibit CD4^+^ T cells, including T_FH_ function and differentiation, during vaccination responses and in vitro(41, 43, 95), further suggesting a role of the hypoxia/adenosine axis in GC immunosenescence. Importantly, hypoxic conditions initiate HIF-1α-dependent A2bR expression (96), and adenosine signaling through A2a and A2b receptors upregulates CXCR4 for CXCL12-dependent chemotaxis in multiple cell types(97, 98). Hypoxia-induced HIF-1α can directly upregulate CXCR4 surface expression(99) while repressing GC T_FH_ differentiation and function, likely enhancing GC and T_FH_ dysfunction in aging(100). Simultaneously, signaling through A2aR increases intracellular cAMP which has been linked to elevated CXCR4 expression on T lymphocytes(101). Taken together, we propose a model in which age-associated dysregulation of the hypoxia/adenosine axis drives immunosenescent GC responses through elevated adenosine/A2aR/A2bR-induced CXCR4 expression on GC T_FH_ cells (Figure 5E). Accordingly, we believe pADA co-immunization may, in turn, restore aged GC T_FH_ localization and function by reducing CXCR4 expression through ADA-mediated adenosine metabolism.

Supporting this proposed mechanism, we have demonstrated previously unidentified age-related alterations to adenosine levels in dLNs following immunization and infection (Figure 6F-H). Remarkably, pADA significantly reduced adenosine in aged dLNs compared to pRBD-immunized mice (Figure 6H), which coincided with significantly larger dLNs (Figure 6I). Taken together, these data indicate that aging is characterized by impaired, antigen-driven regulation of adenosine in dLNs, resulting in sustained suppressive signaling during recall responses. pADA co-immunization corrected this defect, coinciding with restored T_FH_ activation (Figure 4) and functional antibody responses (Figure 3), improved GC organization (Figure 5), and enhanced protection during infection (Figure 6). While these measurements do not resolve cell-specific sources of adenosine or downstream receptor signaling, they provide in vivo evidence that ADA’s enzymatic activity counters age-associated immunosenescence by re-establishing appropriate metabolic regulation of the lymph node microenvironment. Importantly, young mice modeling optimal pRBD-induced immunity exhibited superior immunological responses and protection against challenge irrespective of pADA inclusion, supporting aging-specific modulation.

Collectively, these findings position the ADA/adenosine axis as a mechanistically defined and therapeutically actionable pathway for overcoming age-associated T cell and humoral immunosenescence, warranting further investigation to refine its translational potential in aging and vaccine responsiveness. Future work investigating temporal adenosine kinetics, in situ T_FH_ localization, and adenosine signaling pathways within GCs could further elucidate these age-related GC alterations and the modulatory mechanism of ADA.

## Author Contributions

M.A.K., E.K.H., M.C.A., E.N.K., and M.R.B. conceived the original hypothesis and experimental framework for the study. M.A.K., E.K.H., M.C.A., and E.N.K. refined the study design and directed experimental execution. E.N.K. performed immunizations, bleeds, tissue harvests, and immunological assays; coordinated tetramer staining; and analyzed the data. M.C.A. provided the MHC class II tetramer probe, Clostridioides difficile spores, and anaerobic equipment required for challenge studies. D.R.J. performed immunizations, bleeds, and tissue harvests. J.R.M. performed lymphoid organ processing and tetramer-based immunological assays. K.W., and A.O.E. assisted with tissue processing during harvests. C.S. assisted with procurement and coordination of aged mouse cohorts and provided consultation to aging animal model design. M.G.A. provided fluorescent probes and contributed to optimization of B cell staining and analysis.

The manuscript was written by E.N.K. with intellectual guidance and critical review from M.A.K., E.K.H., and M.C.A. All authors approved the final manuscript. E.N.K. and M.A.K. coordinated manuscript submission, with M.A.K. serving as corresponding author.

## Acknowledgments.

We would like to thank the Drexel University College of Medicine Flow Cytometry Core for providing access to BD LSR equipment. Furthermore, we would like to thank Michael C. Abt for assistance in challenge studies and tetramer staining optimization, and for providing spores, an anaerobic chamber, and supplies for challenge studies. We thank the NIH Tetramer Core Facility (NIH Contract 75N93020D00005 and RRID:SCR_026557) for providing the TcdB1961 tetramer and the CLIP control tetramer utilized within this manuscript. Thank you to Yi-Kan Pan and Mohamad-Gabriel Alameh for preparing and providing fluorescent probes for B cell analyses. We would also like to thank Jeffrey R. Maslanka, Nonto Mdluli, Brijesh Karanam, and Minh Tranha for assistance with neutralization assay, tetramer staining, and challenge study optimizations. We thank Matthew Bell for initiating early proof of concept aging studies with the Clostridioides difficile vaccine model, which helped establish the feasibility of investigating vaccine immunosenescence in this system. Finally, we thank the Drexel animal facility for caring for the mice used in these studies and Inovio Pharmaceuticals for use of the CELLECTRA™ device for in vivo electroporation.

## Supplemental Figure Legends

**Supplementary Figure 1.** Young (6-8 weeks) and aged (72-76 weeks) mice were vaccinated with pVax (20ug), pRBDA+pRBDB (5ug/plasmid), or pRBDA+pRBDB+pADA (5ug/RBD plasmid + 10ug pADA) over 28 days. Spleens were harvested at ten days post-boost and processed for ELISpot and flow cytometry. (A) ICS gating strategy and (B) representative plots. (C) CD44+CD62L-CD4 T effector memory cells per 10^6^ splenocytes and (D) Bcl-6+CXCR5^Hi^PD1^Hi^ T_FH_ cells per 10^6^ splenocytes quantified with flow cytometry. (E) Gating strategy for tetramer staining based on the following parameters: Singlets, Live, Lymphocytes, CD3/5+, CD4+ CD44+. Data from three independent experiments were pooled for statistical analysis. Young pVax (n=8), Aged pVax (n=4), Young pRBD (n=15), Aged pRBD (n=10-13), Young pRBD+pADA (n=14-15), Aged pRBD+pADA (n=9-12). (C, D) Error bars indicate mean and SEM. *P<0.05, **P<0.01, ***P<0.001 and ****P<0.0001 by 2-way ANOVA with Fisher’s LSD post-hoc test.

**Supplementary Figure 2.** Frozen splenocytes from young (6-8 weeks) and aged (72-76 weeks) mice vaccinated with pVax (20ug), pRBDA+pRBDB (5ug/plasmid), or pRBDA+pRBDB+pADA (5ug/RBD plasmid + 10ug pADA) over 28 days were thawed and analyzed for RBD-specific B cell responses with flow cytometry. (A) Gating strategy for toxin A and B probe staining based on the following parameters: Lymphocytes, Singlets, Live, CD3-, CD19+, IgM-IgD-. (B) Toxin A-and (C) toxin B-specific probe+ switched B cells per 10^6^ splenocytes. Data from three independent experiments were pooled for statistical analysis. Young pVax (n=6), Aged pVax (n=3), Young pRBD (n=8), Aged pRBD (n=8), Young pRBD+pADA (n=6), Aged pRBD+pADA (n=10). (B, C) Error bars indicate mean and SEM. *P<0.05, **P<0.01, ***P<0.001 and ****P<0.0001 by 2-way ANOVA with Fisher’s LSD post-hoc test.

**Supplementary Figure 3.** (A) Gating strategy for AIM+ T_FH_ analyses based on the following parameters: Lymphocytes, Singlets, Live, CD3+, CD4+, CXCR5+. Frequency and mean fluorescence intensity of (B, C) CD25, OX40, and (B, D) PD-L1 AIM markers on T_FH_ cells. pVax n=1-2/aged group, RBD n=3-7/age group, RBD+ADA n= 7-8/age group. Data from two independent experiments were pooled for statistical analysis. Young pVax (n=2), Aged pVax (n=2), Young pRBD (n=3-8), Aged pRBD (n=7), Young pRBD+pADA (n=8), Aged pRBD+pADA (n=8). (C-D) Error bars indicate mean and SEM. *P<0.05, **P<0.01, ***P<0.001 and ****P<0.0001 by 2-way ANOVA with Fisher’s LSD post-hoc test.

**Supplementary Figure 4.** (A) GC-Tfh CXCR4 expression gating strategy based on the following parameters: Lymphocytes, Singlets, Live, CD3+, CD4+CD8-, CD44+CD62L-, PD-1^Hi^CXCR5^Hi^.(B) Concatenated flow plots of CXCR4+ TcdB_1961_-specific GC T_FH_ (n=5-6/ group). (C) Frequency of TcdB_1961_+ GC T_FH_. (D) Quantification of TcdB_1961_+ GC T_FH_ cells per 10^6^ lymphocytes. (E) Frequency of CXCR4+ TcdB_1961_-specific GC T_FH_. Error bars indicate mean and SEM. Data from two independent experiments were pooled for statistical analysis. Young pRBD (n=6), Aged pRBD (n=6-7), Young pRBD+pADA (n=5-6), Aged pRBD+pADA (n=7). (C-E) Error bars indicate mean and SEM. *P<0.05, **P<0.01, ***P<0.001 and ****P<0.0001 by 2-way ANOVA with Fisher’s LSD post-hoc test.

**Supplementary Figure 5.** Young (6-8 weeks) and aged (72-76 weeks) mice were vaccinated twice with pVax (20ug) (n=6/age group), pRBDA+pRBDB (5ug/plasmid) (n=7/age group), or pRBDA+pRBDB+pADA (5ug/RBD plasmid + 10ug pADA) (n=7/age group) over 28 days, rested for 6 weeks, then challenged with 10^3^ CD196 spores by oral gavage. (A-C) Weight loss and (D-F) morbidity were tracked daily for 13 days post-challenge. Morbidity scores were assigned based on established clinical scoring criteria (see Methods). Error bars indicate mean and SEM. Young pVax (n=6), Aged pVax (n=6), Young pRBD (n=7), Aged pRBD (n=7), Young pRBD+pADA (n=7), Aged pRBD+pADA (n=7). *P<0.05, **P<0.01, ***P<0.001 and ****P<0.0001 by Mixed-Effect Analysis with Geisser-Greenhouse Correction.

