## Supplementary figures and images for "Adenosine deaminase co-immunization reverses age-associated immunosenescence by restoring germinal center T follicular helper cell function"

### Supplemental Figures

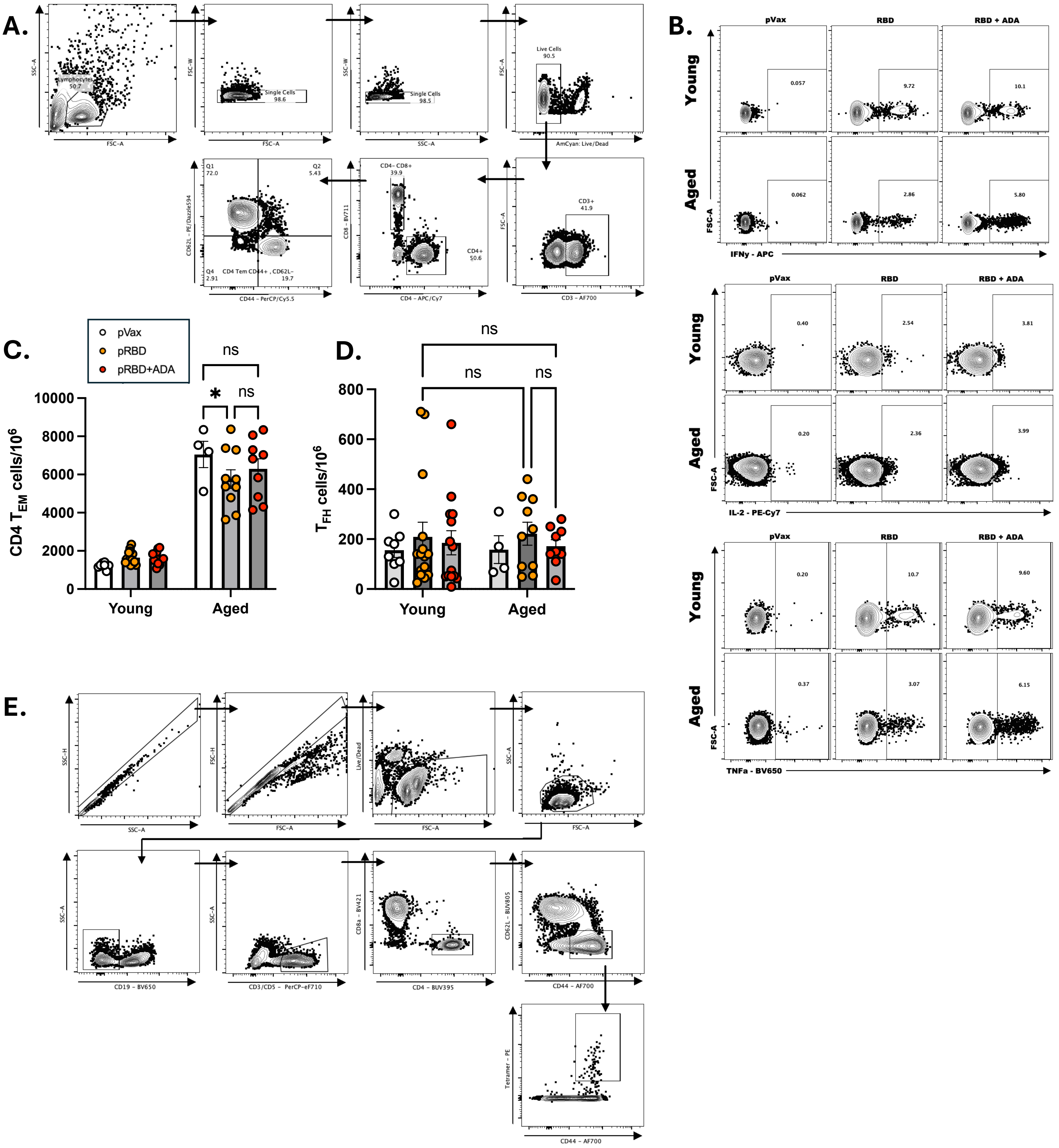

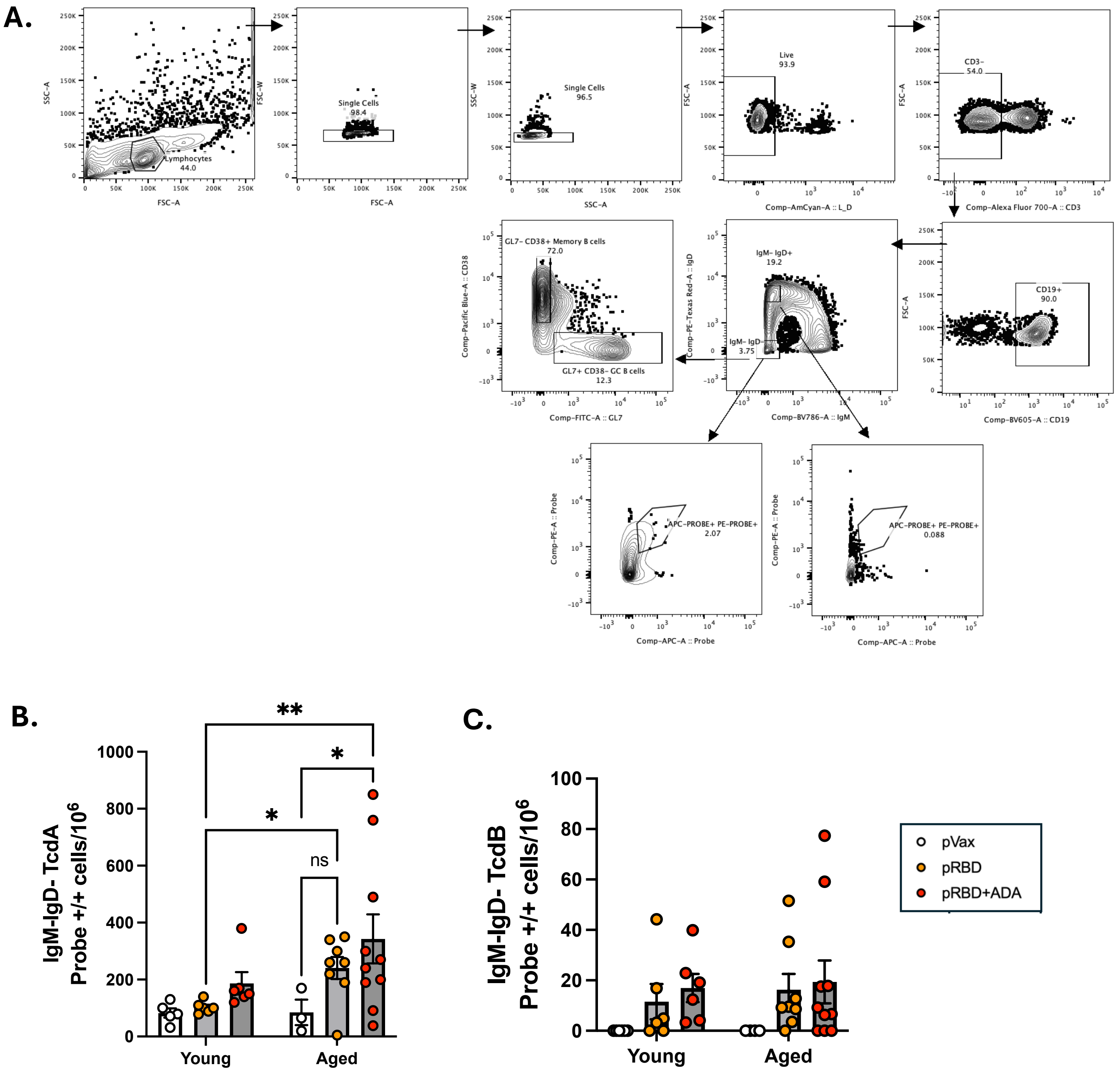

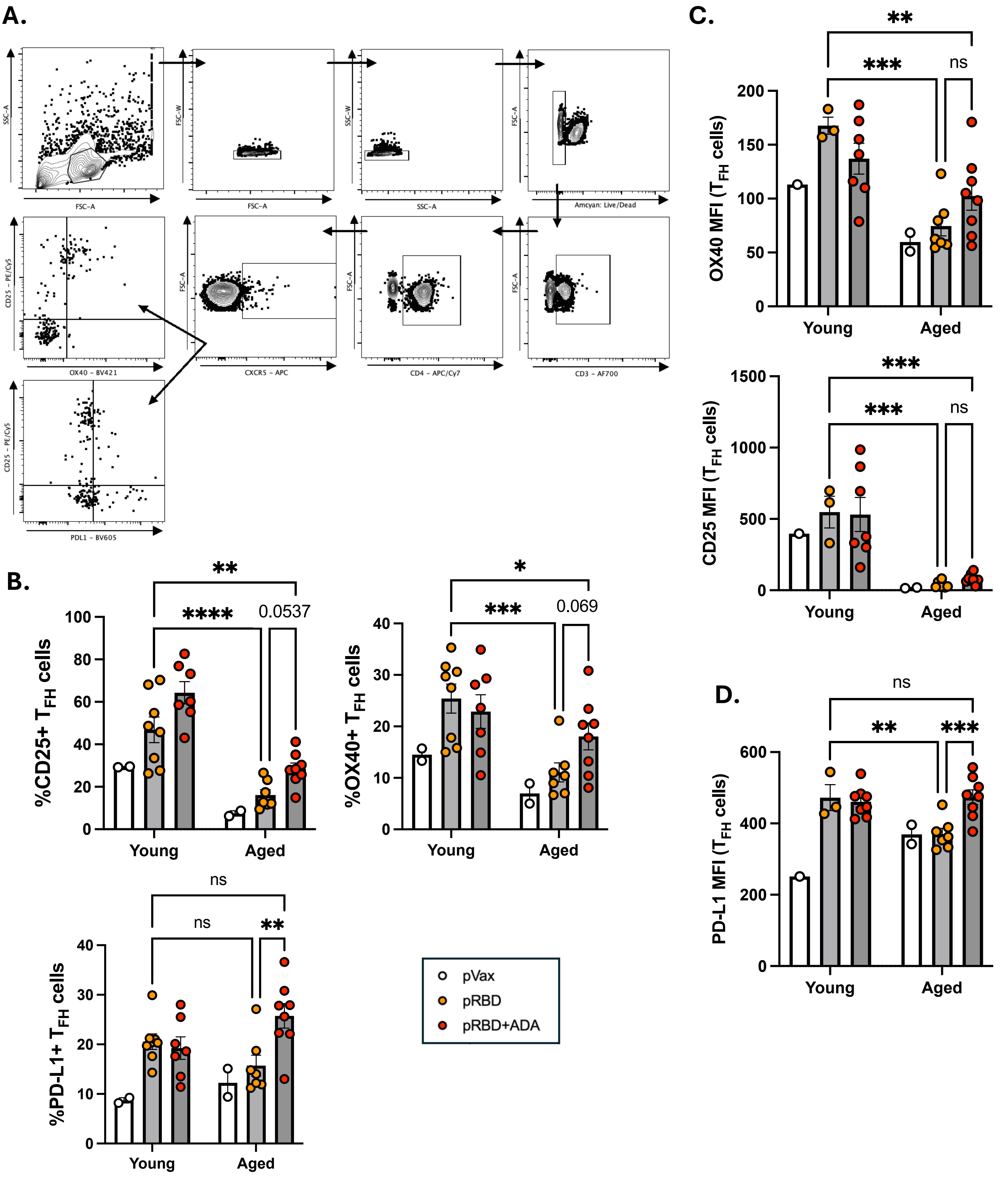

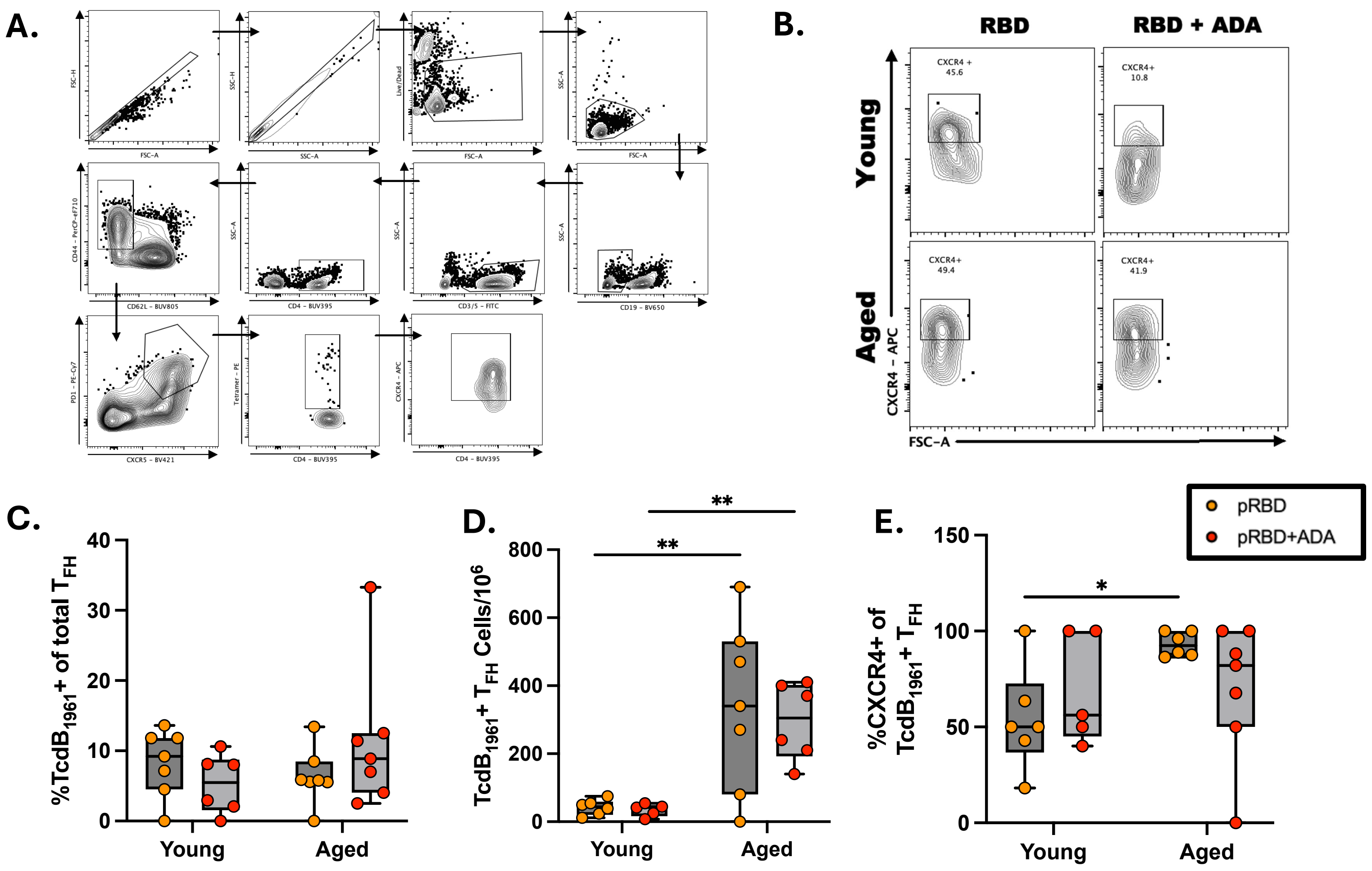

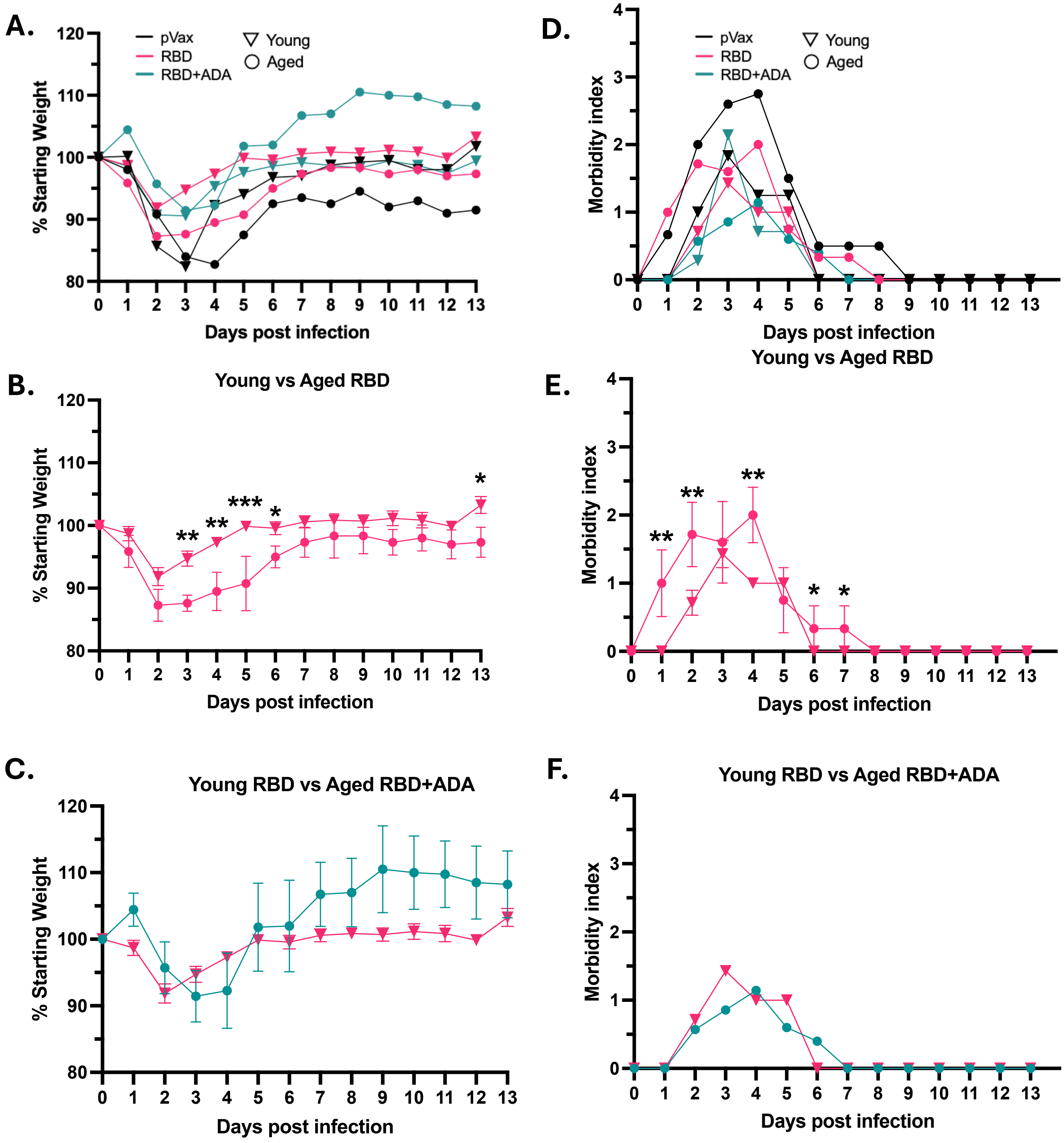
